# Structural and biochemical studies of human PP2A(B55) holoenzyme and ENSA protein complex

**DOI:** 10.64898/2025.12.13.693833

**Authors:** Gang Tian, Xiang Zhang, Hongyue Li, Gangyu Sun, Lulu Xue, Yufeng Yang, Qiyu Mao, Yan Gao, Lei Sun, Guoliang Xu, Zhizhi Wang, Wenqing Xu

**Affiliations:** School of Life Science and Technology, ShanghaiTech University, Shanghai, China; CAS Key Laboratory of Epigenetic Regulation and Intervention, Shanghai Key Laboratory of Molecular Andrology, Shanghai Institute of Biochemistry and Cell Biology, Center for Excellence in Molecular Cell Science, University of Chinese Academy of Sciences, Chinese Academy of Sciences, Shanghai 200031, China; Zhongshan Institute for Drug Discovery, Shanghai Institute of Materia Medica, Chinese Academy of Sciences, Zhongshan 528400, China; Shanghai Fifth People’s Hospital, Shanghai Institute of Infectious Disease and Biosecurity, Shanghai Key Laboratory of Medical Epigenetics and Institutes of Biomedical Sciences, Fudan University, Shanghai 200032, China; Shanghai Institute for Advanced Immunochemical Studies, ShanghaiTech University; Shanghai, 201210, China

**Keywords:** PP2A(B55) holoenzyme, ENSA, SliM, cryo-EM

## Abstract

The serine/threonine protein phosphatase 2A (B55) holoenzyme (PP2A(B55)), a key cell cycle regulator, is tightly regulated by ENSA, whose phosphorylation at Ser67 inhibits PP2A(B55). The structural basis for how pENSA orchestrates this inhibition to govern mitotic progression remains unclear. In this study, we disapproved the previous hypothesis of direct ENSA-PP2A A subunit interaction, and showed that the ENSA-PP2A(B55) interaction requires the PP2A holoenzyme. We then determined cryo-EM structure of the PP2A(B55)-ENSA^S67D^ complex at 3.03 Å resolution, which reveals four distinct regions of ENSA, each engaging specific sites on the B55α or Cα subunits. The structure reveals an extended, inhibited conformation of the PP2A(B55) holoenzyme, which contrasts sharply with the contracted, active state of apo PP2A(B55). pENSA binding induces the C subunit to shift away from the B55 subunit, with the pS67-containing motif inserting into the catalytic site of C subunit and locking the PP2A(B55) holoenzyme in an inactive state. Our work also revealed novel ligand-binding surfaces on B55, and expanded definition of the short linear motif (SliM). Overall, our work provided novel insights into PP2A regulation during cell cycle, and offers new avenues for the discovery of specific PP2A(B55) inhibitors and modulators.

## Introduction

The serine/threonine protein phosphatase 2A (PP2A) plays a vital role in cell signaling, metabolism, cell cycle regulation, and cancer development (1). PP2A accounts for 1% of the total cellular protein and contributes to the main serine/threonine-specific phosphatase activity in mammalian cells. The regulatory B subunit of the PP2A holoenzyme is responsible for the specific recognition of substrates and can be divided into four subfamilies: B (B55, also known as PR55), B’ (B56, also known as PR61), B’’ (PR48, PR59, PR70, PR72, PR130 and G5PR) and B’’’ (PR93, PR110 and zincin) (2).

Alpha-endosulfine (ENSA) was originally identified in sheep brain as an endogenous regulatory ligand of ATP-dependent potassium channels (3). ENSA is expressed in a variety of tissues, suggesting that it has multiple biological functions. ENSA is highly expressed in muscle and brain and low in the pancreas, similar to the tissue distribution of ATP-sensitive K-channels (K-ATP) (4–7). *In vitro* experiments have shown that ENSA can interact with K-ATP channels to regulate insulin release (4). ENSA expression in brain tissue of patients with Alzheimer’s disease and Down syndrome is lower than normal (5, 8).

To initiate cell mitosis, activated Greatwall kinase (GWL) phosphorylates its substrates, ENSA and the related cAMP regulated phosphoprotein-19 (ARPP19), at Ser67 and Ser62, respectively. These phosphorylated proteins (pENSA/pARPP19) then bind to and potently inhibit the PP2A(B55) holoenzyme. The resulting suppression of PP2A(B55) activity prevents the dephosphorylation of crucial CDK1 regulators, such as Wee1 and Cdc25, thereby ensuring the hyperactivation of the Cyclin B/CDK1 complex and driving mitotic entry. Conversely, termination of mitosis is contingent upon the inactivation of this inhibitory loop (9). The PP2A(B55) holoenzyme itself catalyzes the dephosphorylation of its bound inhibitors, pENSA (10) and pARPP19 (11). This auto-deinactivation mechanism leads to the reactivation of the phosphatase, ultimately mediating the transition out of mitosis (9).

ENSA and ARPP19 protein sequences are highly conserved, particularly at residues S67/S62 (12), suggesting similar biological roles. However, important differences exist – ENSA has more negatively charged amino acids at its N-terminus than ARPP19 (13), which may alter how each interacts with PP2A. Notably, only ENSA can regulate the transcription factor Treslin via the PP2A(B55) holoenzyme during Cyclin B/CDK1-activated mitosis, thereby influencing the S phase (14).

Conflicting models have been proposed for the mechanism of ENSA-mediated inhibition of the PP2A(B55α) holoenzyme (13). A previous study by Thapa *et al*. reported a direct interaction between wild-type ENSA (ENSA^WT^) and the isolated Aα subunit, measuring a dissociation constant (*K*_D_) of 3.9 µM via Microscale thermophoresis (MST) (13). Similarly, the same group also reported a direct interaction between wild-type ARPP19 (ARPP19^WT^) and the isolated Aα subunit, with a measured *K*_D_ of 7.9 µM via MST (15). Based on this result and other biochemical data, they proposed a model in which ENSA and ARPP19 inhibit the PP2A(B55α) holoenzyme primarily by engaging the scaffolding Aα subunit, rather than the regulatory (B55) or catalytic (Cα) subunits (13, 15). However, recent structural studies of the PP2A(B55α)-tpARPP19 (thiophosphorylated ARPP19) complex demonstrated direct interactions of ARPP19 with the B55 and Cα subunits, instead of the Aα subunit (16).

It has been reported that both regulators and substrates of the PP2A(B56) holoenzyme interact with the B56 subunit through a conserved ‘SliM motif’ (LxxIxE) (17–20). While the interaction mechanism for the PP2A(B56) holoenzyme is relatively well understood, the mechanisms governing binders of the PP2A(B55) holoenzyme (including inhibitors, recruiters, and substrates) are less clear. Recently reported structures of PP2A(B55α) binders include those of inhibitors (tpARPP19, PDB ID 8TTB (16); FAM122A, PDB ID 8SO0 (16); and B55i, PDB ID 9N0Z (21)), recruiters (Eya3, PDB ID 9N0Y (21), 9C7T (22); and IER5, PDB ID 8UO5 (23)), and the substrate p107 (PDB ID 9C6B (22)). A key question is whether ENSA shares regulatory mechanisms similar to these diverse binders.

In this study, we assembled the PP2A(B55)-ENSA^S67D^ (the phosphomimicking mutant) quaternary complex and determined the high resolution structure using single-particle cryo-EM. Combined with biochemical data, we provide new insights for the allosteric regulation of the PP2A(B55) holoenzyme by pENSA and other binders.

## Results

### ENSA does not directly interact with the PP2A Aα subunit

Thapa *et al*. reported a direct interaction between wild-type ENSA (ENSA^WT^) and the Aα subunit (13). In GST-pull assays, we found that neither GST-tagged ENSA^WT^ nor the phosphomimetic mutant ENSA^S67D^ interacted with the isolated Aα or B55α subunits, or with the AC core enzyme. In contrast, robust binding was observed only with the intact PP2A(B55α) holoenzyme, with ENSA^S67D^ exhibiting significantly stronger binding than ENSA^WT^ (Figure 1A, S1A). In parallel, our MST experiments, revealed a very weak binding between the Aα subunit and ENSA (with *K*_D_ with ENSA^WT^ and ENSA^S67D^ of ∼600 and 800 µM, respectively), which is unlikely to be physiologically relevant (Figure 1B). This lack of direct interaction was further confirmed using two orthogonal biophysical methods, Isothermal titration calorimetry (ITC) and Biolayer interferometry (BLI), both of which showed essentially no binding between ENSA^WT^, or ENSA^S67D^, and the isolated Aα subunit (Figure 1C-F, Table S1). In contrast, the binding affinity between PP2A A-B55-C holoenzyme and ENSA^WT^ was measured to be 1.45 μM (Figure 1G, Table S1).

**Figure 1.**
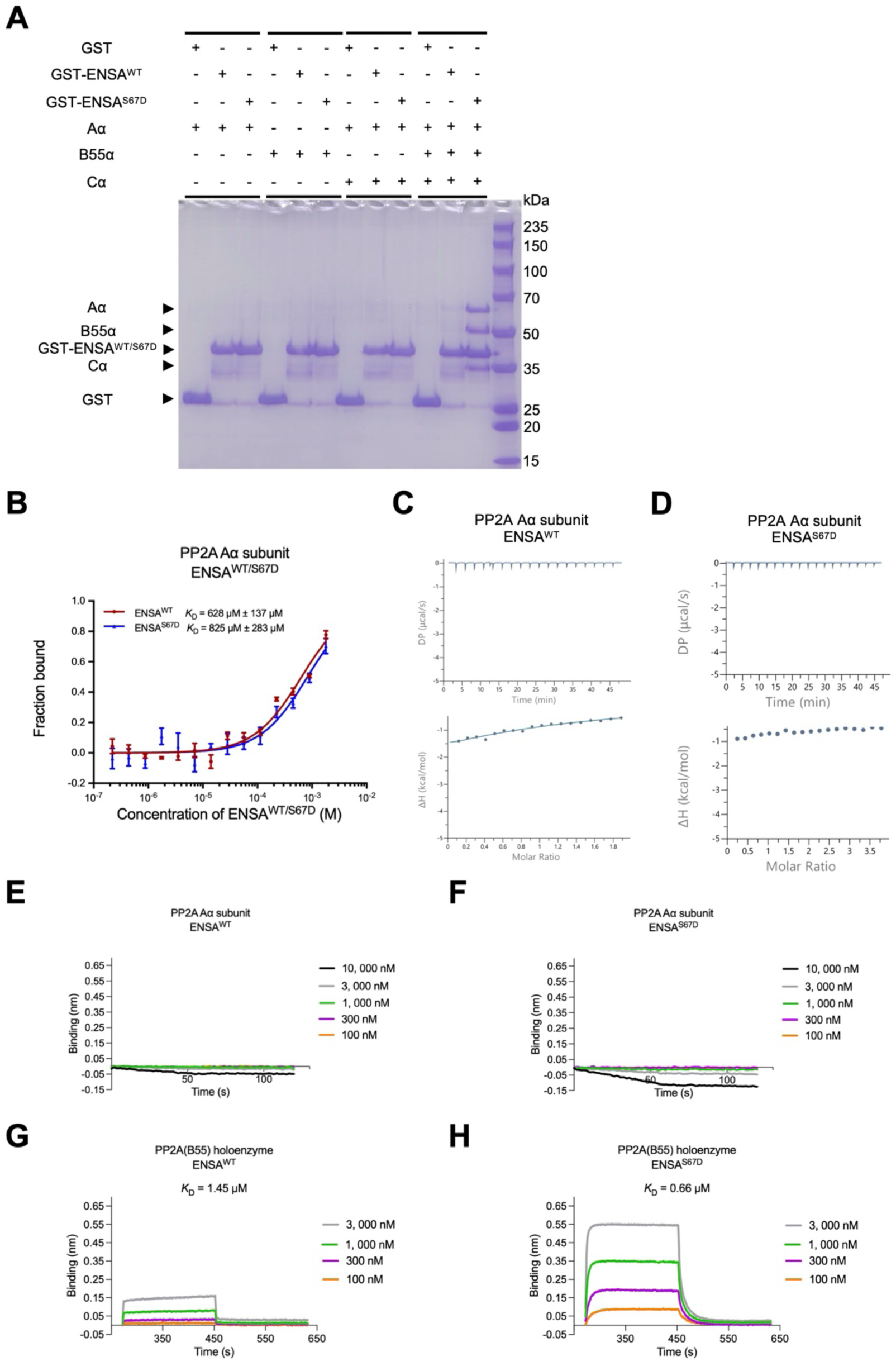
ENSA interacts stronger to PP2A(B55) holoenzyme that the A subunit. **A.** GST pull-down assays showing the interactions of wild-type ENSA (ENSA^WT^) or the phosphomimetic mutant ENSA^S67D^ with the PP2A(B55α) Aα subunit, B55α subunit, AC core enzyme, and intact holoenzyme. **B.** Microscale thermophoresis (MST) analysis of the interactions between ENSA^WT/S67D^ and the PP2A Aα subunit. **C** and **D**. Isothermal titration calorimetry (ITC) analysis of the interactions between the Aα subunit and ENSA^WT^ (C) or ENSA^S67D^ (D). **E**-**H**. Biolayer interferometry (BLI) analysis of the interactions of ENSA^WT^ (E, G) and ENSA^S67D^ (F, H) with the Aα subunit (E, F) or the intact PP2A(B55α) holoenzyme (G, H). Dissociation constants (*K*_D_) calculated from these data are summarized in Table S1. Details of data processing are provided in the Methods.

To measure how phosphorylation affects ENSA’s binding, we used BLI. Our results showed that the phosphomimetic mutant ENSA^S67D^ binds to the PP2A(B55α) holoenzyme with a *K*_D_ of 0.66 µM, which is significantly stronger than the affinity seen with ENSA^WT^ (with a *K*_D_ of 1.45 µM; Figure 1G, H, Table S1). We therefore conclude that ENSA’s binding is dependent on the fully assembled PP2A(B55α) holoenzyme, and ENSA S67 phosphorylation enhances this interaction.

### ENSA^S67D^ binding to the PP2A(B55α) holoenzyme via dual subunit contacts

PP2A(B55α) holoenzyme is formed by interaction of the B55α regulatory subunit to the PP2A-AC core complex. We next analyzed the binding of ENSA to PP2A A and C subunits. The direct binding of both ENSA^WT^ and ENSA^S67D^ to the isolated B55α subunit was almost undetectable in our GST pull-down assays (Figure 1A, S1A), while showed weak affinity in BLI assays (Figure S1B, C, Table S1). In contrast, interaction with the Cα subunit was strictly phosphorylation-dependent, as only ENSA^S67D^ exhibited readily detectable binding signal (Figure S1D, E, Table S1). Consistent with the requirement for B55α, neither ENSA variant bound to the PP2A-AC core enzyme (Figure S1F, G, Table S1).

Thus, high-affinity binding of ENSA to PP2A requires both the B55α subunit as a primary docking site and a phosphorylation-dependent secondary contact with the Cα subunit, suggesting a multi-faceted binding and regulatory mechanism.

### Overall cryo-EM structure of PP2A(B55α)-ENSA^S67D^ complex

To elucidate the molecular mechanism of ENSA-mediated inhibition, we determined the structure of the PP2A(B55α) holoenzyme in complex with the phosphomimetic mutant ENSA^S67D^ using single-particle cryo-EM. The stable quaternary complex was purified by size-exclusion chromatography (SEC), and the peak fraction from SEC was collected, supplemented with additional ENSA^S67D^ at a 5-fold molar excess, and used to prepare cryo-EM samples (Figure S2A). Cryo-EM micrographs displayed a high density of well-distributed particles (Figure S2B). Following iterative 2D and 3D classification, mask, and EMReady (24) methods, a final dataset of 227,045 particles yielded a high-resolution reconstruction of the complex with an overall resolution of 3.03 Å (Figure S3).

The structure reveals an extensive interface between ENSA^S67D^ and the PP2A(B55α) holoenzyme, mediated by three distinct helices (Figure 2A, B). Helices 1 and 2 of ENSA^S67D^ wrap around the top and side of the B55α subunit, respectively, acting as a primary docking elements. The helix 3 extends towards the Cα subunit, with its D67 residue—mimicking the critical pS67—inserted directly into the phosphatase catalytic site. The overall interaction spans four distinct regions of ENSA, each engaging specific sites on the B55α or Cα subunits (Figure 3A, S4A).

**Figure 2.**
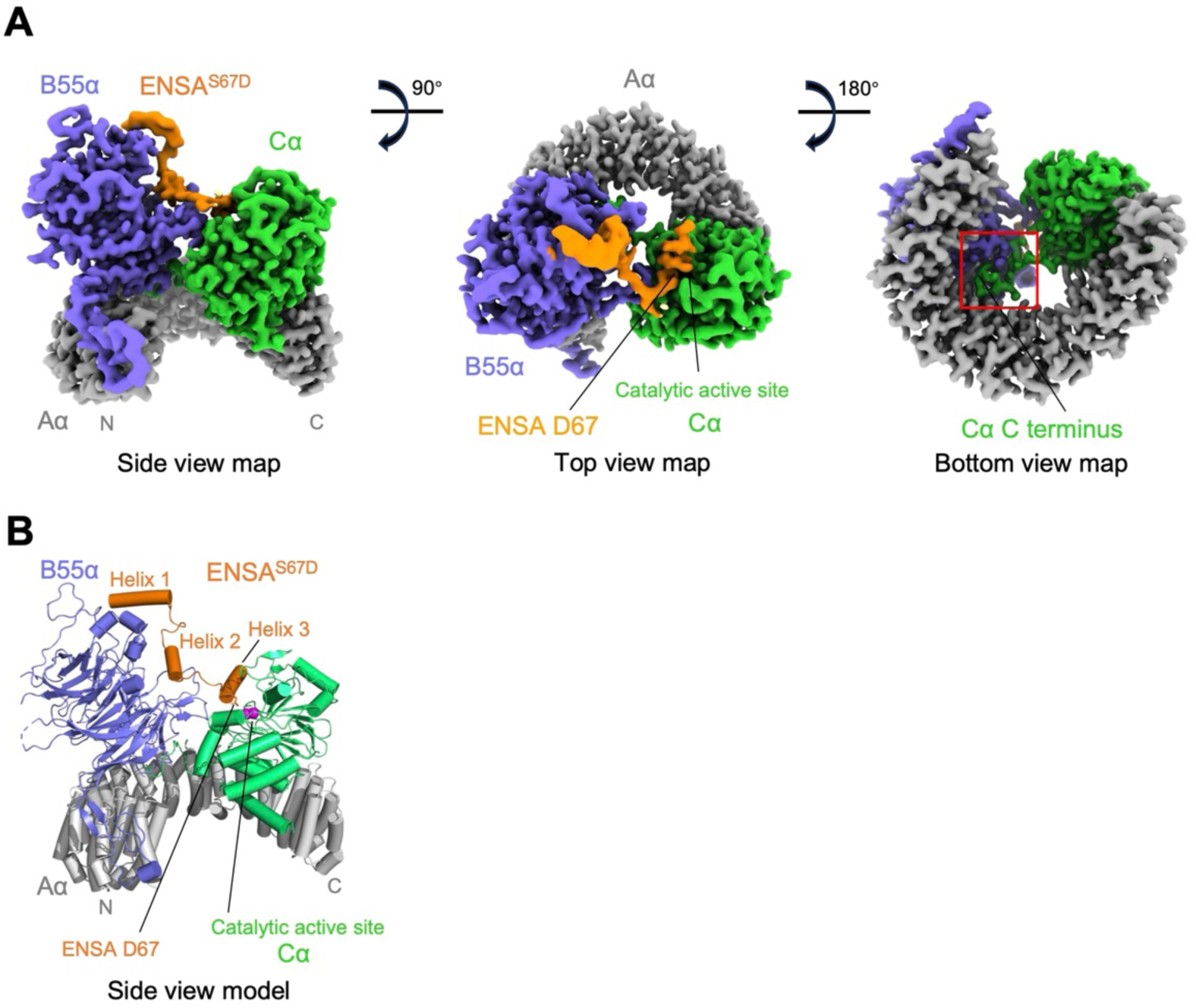
Cryo-EM structure of the PP2A(B55)-ENSA^S67D^ complex. **A.** Side, top, and bottom views of the cryo-EM density map of the PP2A(B55)-ENSA^S67D^ complex (this study, PDB ID 9XGY, under release), color-coded by subunit. The Aα, B55α, and Cα subunits, and ENSA^S67D^ are colored dark gray, medium slate blue, lime green, and dark orange, respectively. **B.** Side view of the complex in cartoon representation. The key residue D67 of ENSA is shown as sticks. The Aα, B55α, Cα, and ENSA^S67D^ are colored gray, slate, lime green, and orange, respectively.

**Figure 3.**
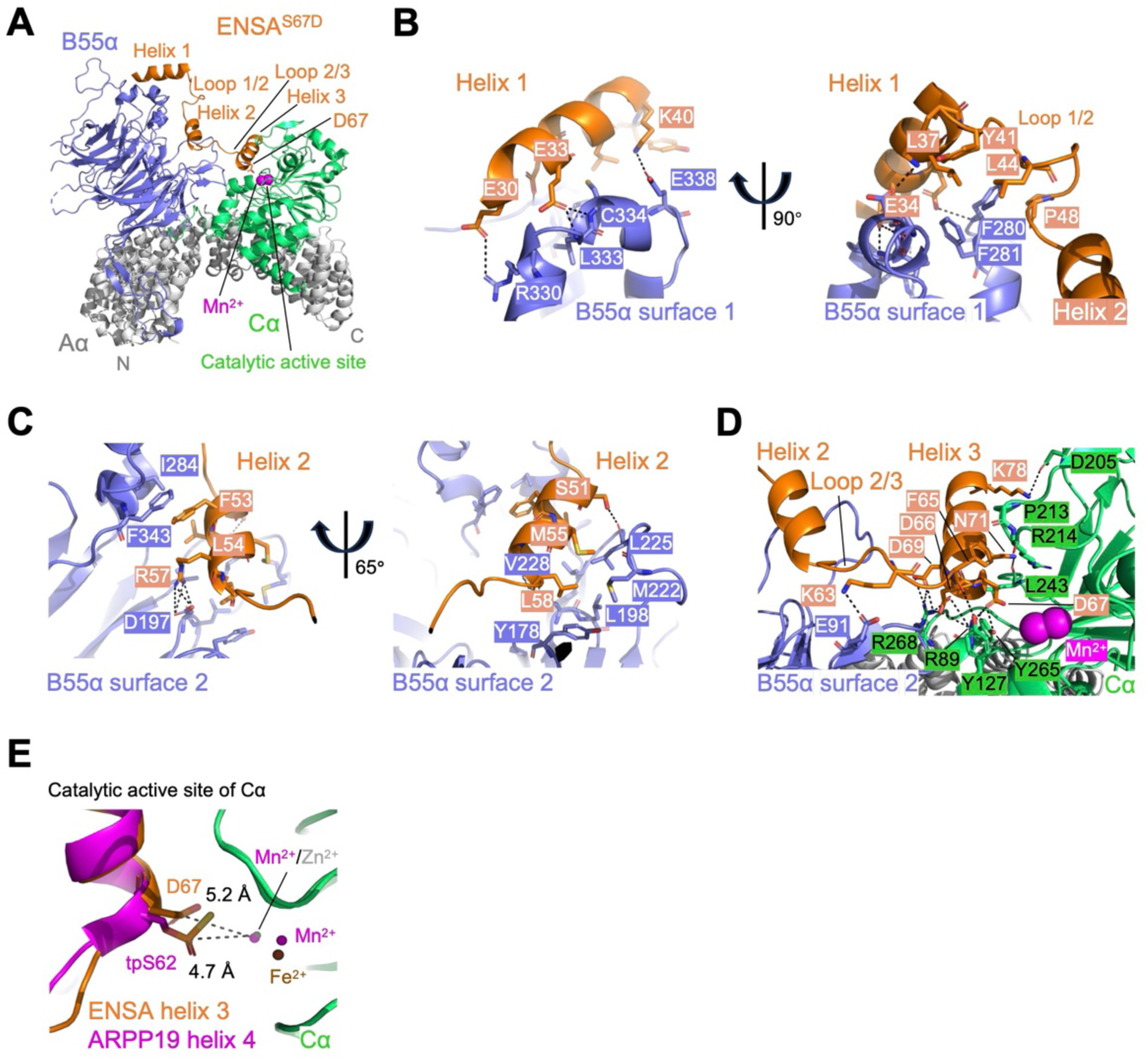
Detailed interfaces between ENSA^S67D^ and PP2A(B55) holoenzyme. **A.** Overview of the complex in cartoon representation. The subunits are colored as in Figure 2B. **B.** Close-up view of the ENSA helix 1 and its adjacent loop with the top of the B55α subunit. Key interacting residues are shown as sticks. **C.** Detailed view of the ENSA short linear motif (SliM) (Helix 2) binding to the conserved Surface 2 pocket on B55α. **D.** The interface where ENSA Loop 2/3 and helix 3 contact both B55α and the catalytic Cα subunit, bridging the two core components of the holoenzyme. **E.** Comparison of the catalytic site engagement. Distances are shown from the key inhibitory residue (D67 of ENSA or tpS62 of ARPP19) to the catalytic metal ions (Mn²⁺ in our structure; Zn²⁺ in PDB ID 8TTB). Fe^2+^, Mn^2+^, and Zn^2+^ are shown as spheres scaled to 0.2. **D** and **E**. Fe^2+^, Mn^2+^, and Zn^2+^ are colored brown, magenta, and gray (50%), respectively.

### Structural basis for the ENSA^S67D^-PP2A(B55) interaction

The N-terminal helix 1 of ENSA, along with its adjacent loop (Loop 1/2), forms a primary docking site on the top surface of the B55α seven-blade β-propeller (Figure 3B). The interface is cemented by a hydrophobic patch, where ENSA L37, Y41, L44, and P48 nestle into a pocket on B55α lined by B55α F280 and F281. Alanine substitution of these hydrophobic residues (L37A, Y41A, L44A) significantly weakened the interaction, with an effect comparable to deleting the entire N-terminal region (residues 1-45), highlighting the critical role of this hydrophobic core (Figure 4A). The hydrophobic interactions are buttressed by salt bridges formed on both sides of the helix 1: ENSA E30 with B55α R330; ENSA K40 with B55α E338. Mutation of ENSA E30 or K40 to alanine (E30A, K40A) resulted in a modest but detectable reduction in binding affinity (Figure 4A). In addition, the sidechain of ENSA E33 forms hydrogen bonds with the backbone of L333 and C334 of B55α, while ENSA E34 also forms a hydrogen bond with the backbone of B55α F280.

**Figure 4.**
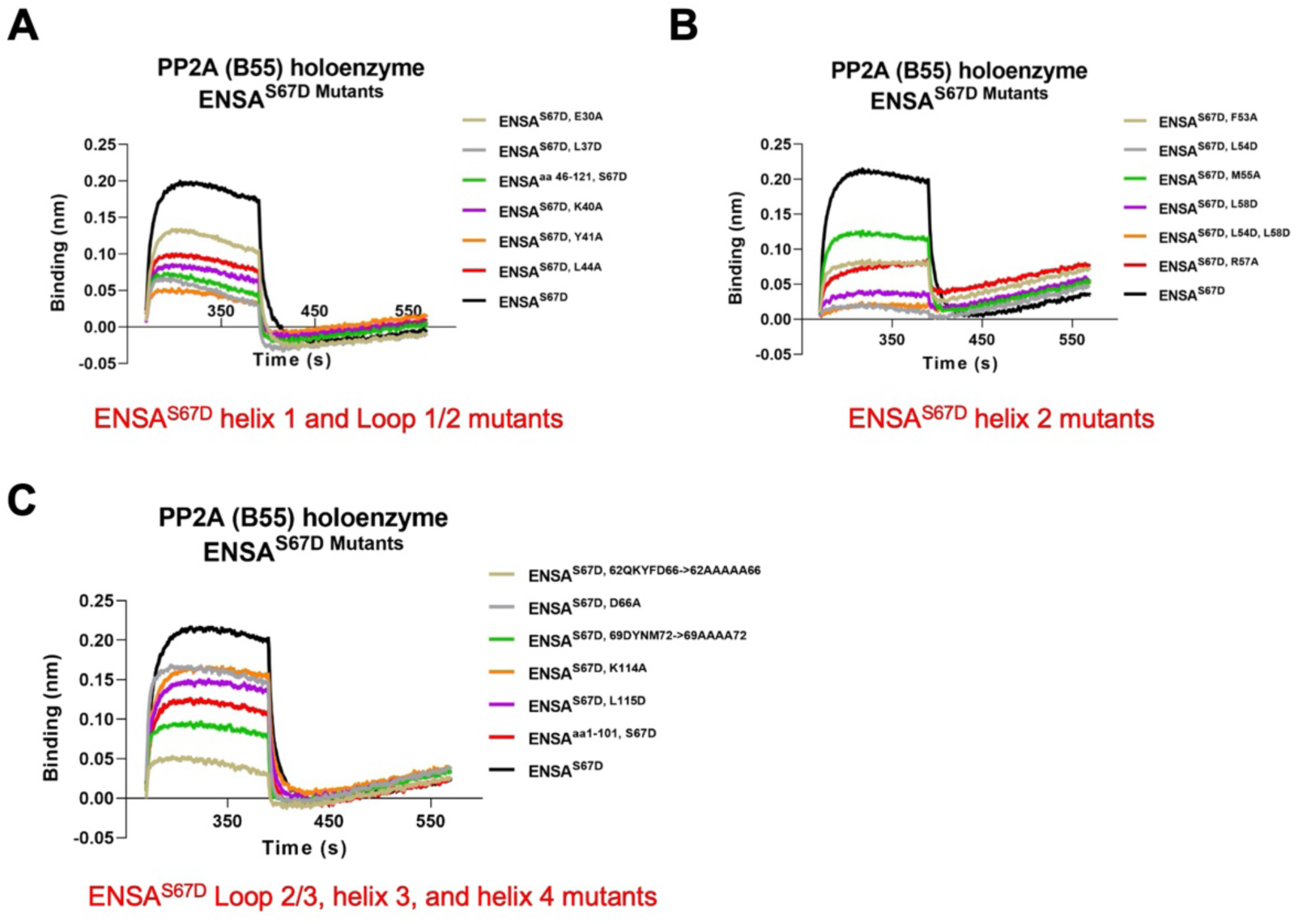
Structure-guided mutants and truncations of ENSA^S67D^ on PP2A interaction using BLI. BLI measuring the binding of various structure-guided ENSA mutants and truncation constructs to the PP2A(B55α) holoenzyme. Details of data processing are provided in the Methods. **A.** Analysis of mutations in ENSA helix 1 and its adjacent loops. **B.** Analysis of mutations within the ENSA SliM (Helix 2). **C.** Analysis of mutations and truncations affecting the C-terminal regions of ENSA (Loop 2/3, helix 3, and helix 4).

Helix 2 and the subsequent loop (Loop 2/3) form the conserved short linear motif (SliM) (16), ⁵³F-L-M-x-R-L-xxxx-K⁶³ in *Homo sapiens* ENSA (*Hs*-ENSA), which engages the Surface 2 on B55α (Figure 3C, D, S5, S6). The SliM interaction is buttressed by two distinct hydrophobic sub-pockets. The first involves ENSA F53 and L54 contacting B55α I284 and F343. The second involves ENSA M55 and L58 interacting with a larger hydrophobic surface on B55α comprising Y178, L198, M222, L225, and V228 of B55α. Mutational analysis underscored the importance of these contacts, with L54D and L58D causing a substantial loss of binding (Figure 4B). Notably, ENSA L54 is the most conserved residue across all ENSA/ARPP19 orthologs, suggesting it forms the foundational hydrophobic anchor for the SliM (Figure S5, S6). Importantly, in this SliM motif, the conserved basic residue ENSA R57 is critical to this interaction, forming hydrogen bonds and salt bridges with B55α D197; the R57A mutation moderately impaired binding (Figure 4B). Additionally, ENSA S51 forms a hydrogen bond with the backbone of B55α L225, while the conserved ENSA K63 forms a salt bridge with B55α E91, further stabilizing the complex (Figure 3D).

Helix 3 of ENSA extends away from B55α to directly engage the Cα catalytic subunit (Figure 3D). The phosphomimetic residue ENSA D67 is positioned deeply within the active site, forming salt bridges with Cα R89, and hydrogen bonds with Cα Y265, respectively. Its carboxylate group is located 5.2 Å from the catalytic metal ions, a position analogous to the thiophosphate group in the tpARPP19-bound structure (4.7 Å) (16), confirming its role as a competitive inhibitor (Figure 3E). The helix is further tethered to Cα by several interactions: ENSA F65 packs against of Cα Y127, ENSA D66 forms hydrogen bonds and salt bridges with R89 and R268 of Cα, ENSA D69 forms salt bridges with Cα R268, ENSA N71 forms a hydrogen bond with the mainchain of Cα L243 and ENSA K78 forms a salt bridge with the Cα D205. Consistent with the importance of this region, a mutant where residues 62-66 were replaced by alanines (62QKYFD66 -> 62AAAAA66) or a 69DYNM72 -> 69AAAA72 mutant both dramatically reduced binding affinity (Figure 4C).

While the C-terminal tail of ENSA (Helix 4) was not visible in our cryo-EM map, suggesting flexibility, its corresponding region in ARPP19 (Helix 5) was resolved in a previous structure (16). Deletion of the ENSA C-terminus residues 102-121 or mutation of two conserved residues within this region, K114A and L115D, significantly impaired binding affinity (Figure 4C). This suggests that this flexible tail may enhance the binding by transiently interacting with the holoenzyme. AlphaFold3 modeling (25) predicts that this tail docks onto B55α, with ENSA K114 forming a salt bridge with B55α D29 and ENSA L115 making hydrophobic contacts with I31, V53, F55, and L439 of B55α (Figure S4A), thus providing an additional, albeit dynamic, layer of interaction.

### The inhibitors engage conserved surfaces on PP2A(B55α) through distinct binding modes

We compared the sequences of the three major classes of PP2A(B55α) inhibitors: ENSA, ARPP19, and FAM122A. While ENSA and ARPP19 are highly conserved across vertebrates (*Hs*, *Mus musculus* (*Mm*), *Gallus gallus* (*Gg*), *Lacerta agilis* (*La*), *Xenopus laevis* (*Xl*), and *Danio rerio* (*Dr*)), suggesting a similar mode of interaction in these organisms, their evolutionary paths diverge in lower eukaryotes. ENSA orthologs are present in organisms like *Drosophila melanogaster* (*Dm*), *Caenorhabditis elegans* (*Ce*), and *Schistosoma mansoni* (*Sm*), whereas ARPP19 is absent in these organisms, suggesting that ENSA has a broader phylogenetic distribution than ARPP19 (Figure S5). Notably, the *Ce*-ENSA sequence show significant divergence from their vertebrate counterparts, hinting at potentially different regulatory mechanisms (Figure S5).

To identify functionally important regions, we first performed a conservation analysis on the B55α and Cα subunits. This analysis revealed five highly conserved surfaces on B55α (Figure 5A). Surfaces 1-3 correspond to previously identified interaction surfaces (16), while Surfaces 4 and 5 are novel conserved surfaces discovered in this study. Similarly, a highly conserved patch is located at the catalytic active site of the Cα subunit (Figure S7A). These conserved regions likely represent critical hubs for regulatory interactions.

**Figure 5.**
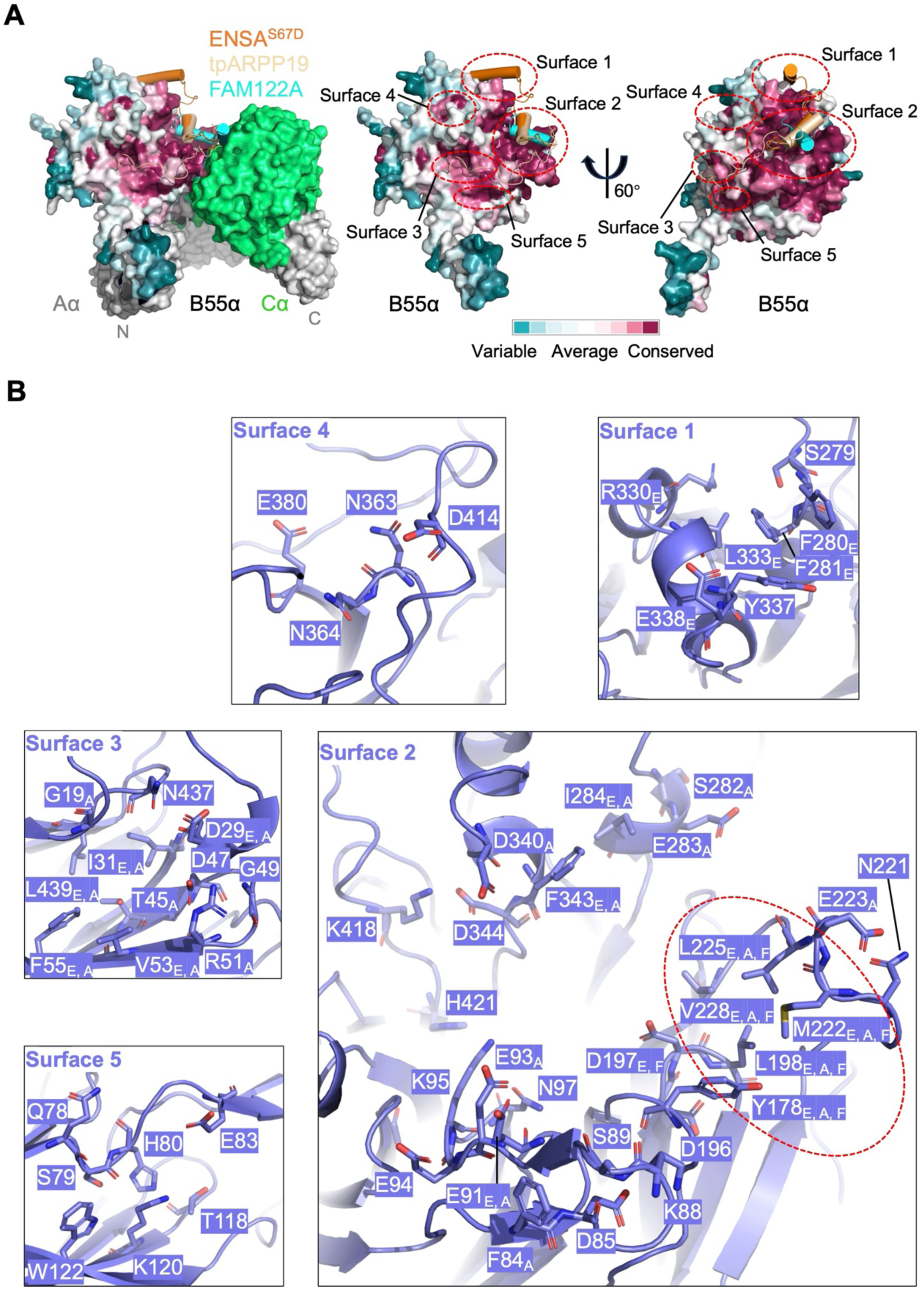
The conserved amino acid analysis of B55α subunit. **A.** The conserved amino acid analysis of the B55α subunit across 150 organisms is generated using the ConSurf server. The B55α subunit is shown as surface, while ENSA, ARPP19 and FAM122A are shown as cartoons. The Surfaces 1-3 are the same as those defined by Padi *et al*. (16), while Surfaces 4 and 5 are newly identified conserved surfaces on B55α. **B.** Base on PP2A(B55)-ENSA^S67D^ (this study), PP2A(B55)-tpARPP19 (thiophosphorylated ARPP19) and PP2A(B55)-FAM122A, the highly conserved functional and structural residues are show as sticks. The residue R330_E_ of B55α specifically interacts with ENSA, while E283_A_ specifically interacts with ARPP19. The L225_E, A, F_ of B55α interacts with ENSA, ARPP19 and FAM122A. The five residues that interact with ENSA, ARPP19, and FAM122A are circled by a red dashed oval.

The three inhibitors—ENSA, ARPP19, and FAM122A—engage the B55α subunit with markedly different footprints (Figure S8-9). Both *Hs*-ENSA and *Hs*-ARPP19 use a multi-site binding mode, contacting Surfaces 1-3 on B55α. In contrast, *Hs*-FAM122A relies on a more focused interaction, engaging only Surface 2. Even within the shared Surface 2, the mode of engagement differs. *Hs*-ENSA (via helix 2) and *Hs*-ARPP19 (via helix 3) dock onto the surface of Surface 2, whereas *Hs*-FAM122A (via helix 1) inserts more deeply into a groove within this surfasce. This results in a distinct helical orientation for FAM122A relative to ENSA and ARPP19 (Figure S8A-C) and a smaller interaction interface (Figure 5, S9). The divergent inhibitors from other species also exhibit unique binding patterns. For instance, the AlphaFold3-predicted model shows that *Ce*-ENSA interacts with Surfaces 1, 2, and the newly identified Surface 5, a binding footprint distinct from that of the human inhibitors (Figure S4B, S8A, B).

While the interactions with B55α are diverse, all inhibitors share a common mechanism for occluding the Cα catalytic site. A key acidic or phosphorylated residue—D67 in *Hs*-ENSA^S67D^, tpS62 in *Hs*-ARPP19, E104 in *Hs*-FAM122A, and pS61 in *Ce*-ENSA—acts as a “plug” by inserting directly into the bi-metal center of the Cα active site (Figure S8B, D). Consequently, the Cα residues that contact these inhibitor “plugs” are highly conserved across all complexes (Figure S7B), consistent with the extreme evolutionary conservation of the Cα subunit itself (Figure S10). However, despite this shared feature, the orientation of the helix presenting the plug differs; for instance, helix 2 of *Hs*-FAM122A has an inverted N-to-C terminal orientation compared to the corresponding helices in ENSA and ARPP19 (Figure S7A, S8B, D), highlighting the remarkable plasticity of inhibitor recognition.

### A conserved pocket on B55α serves as the universal docking site for inhibitor SliM motifs

Our structural analysis reveals that among the five conserved surfaces on the B55α subunit, only Surface 2 serves as the direct binding surface for the SliM present in each inhibitor (Figure 5, S8C). This surface is strategically positioned; it not only constitutes the largest conserved patch on B55α but is also located in close proximity to the catalytic active site of the Cα subunit (Figure 5).

While the core of Surface 2 is critical for SliM recognition, residues at its periphery, such as those adjacent to Surface 4 (e.g., D344, K418, and H421), do not participate in the interaction with any of the inhibitors analyzed (Figure 5). This defines a highly specific and conserved binding pocket for the SliM. Indeed, despite significant sequence divergence, the SliM motifs from phylogenetically distant orthologs, such as *Ce*-ENSA, engage this Surface 2 pocket in a remarkably similar fashion to their human counterparts, *Hs*-ENSA and *Hs*-ARPP19 (Figure S8A-C), establishing Surface 2 as a universal docking hub for this class of inhibitors.

A conserved R (Arginine) residue within the SliM motif has been previously highlighted as a key determinant for interaction with the PP2A(B55α) holoenzyme. Our sequence analysis broadens this definition, revealing that while this position is indeed an arginine in human inhibitors, it is occupied by a K (Lysine) in more distant orthologs such as *Ce*-ENSA (Figure S5, S6). The models predict that these lysine residues (K52 in *Ce*-ENSA helix2) insert into the same pocket within Surface 2 on B55α as the conserved arginines from their human counterparts (*Hs*-ENSA R57, *Hs*-ARPP19 R52, and *Hs*-FAM122A R84) (Figure S4B, S8A, C). Importantly, this conserved basic residue, whether an arginine or a lysine, forms key electrostatic interactions (salt bridges and hydrogen bonds) that anchor the SliM motif to the B55α subunit. This indicates that the presence of a basic side chain at this position, rather than the specific identity of arginine or lysine, is the critical conserved feature for inhibitor recognition.

### Conformational change of PP2A(B55α) induced by ENSA^S67D^ and other binders

To examine the PP2A conformational changes induced by ENSA^S67D^, we compared our structure with recently reported structures of the apo PP2A(B55α) holoenzyme (PDB ID 9MZW (21)) and with those bound to inhibitors (tpARPP19, PDB ID 8TTB (16); FAM122A, PDB ID 8SO0 (16); and B55i, PDB ID 9N0Z (21)), recruiters (Eya3, PDB ID 9N0Y (21), 9C7T (22); and IER5, PDB ID 8UO5

(23)), and the substrate p107 (PDB ID 9C6B (22)). In the apo PP2A(B55α) holoenzyme, the B55α and Cα subunits interact tightly. Cα Y91 forms hydrogen bonds with B55α D209, Cα R89 forms hydrogen bonds with B55α N174, and R214 in the upper lip loop of Cα forms hydrogen bonds with S89 in the β1-β2 hairpin of B55α (Figure 6A, C). The C-terminal tail of Cα (residues 304-309) simultaneously contacts both the Aα and B55α subunits (Figure 6B, D). K416 in HEAT repeat 11 (“H11”) of Aα forms salt bridges with Cα D290 (Figure 6B, E).

**Figure 6.**
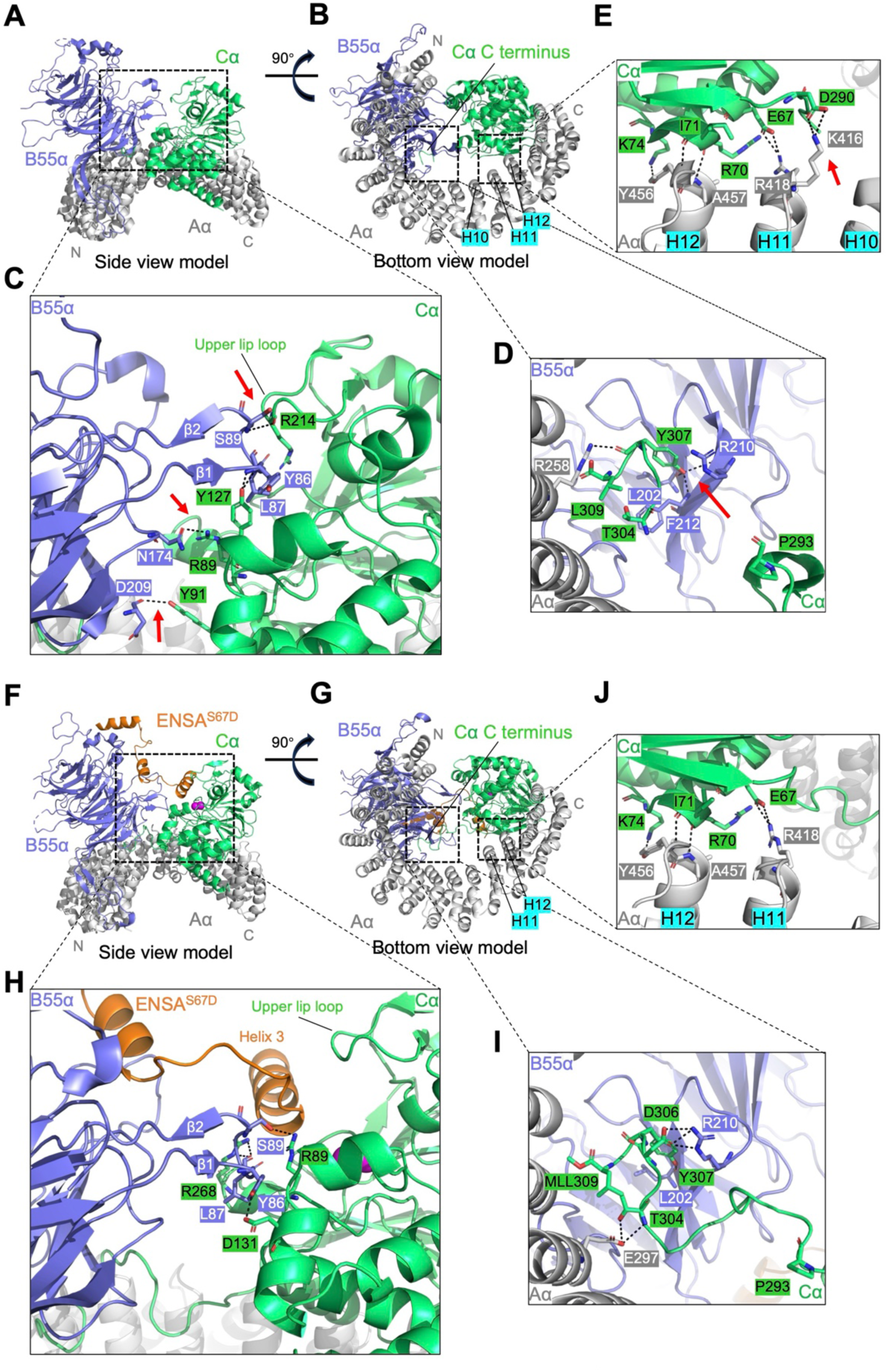
The conformational changes of PP2A upon ENSA^S67D^ binding. **A** and **B**. Side view (A) and bottom view (B) model of apo PP2A(B55) (PDB ID 9MZW). **C.** A zoomed-in view of the interaction between the B55α and Cα subunits from panel (A). **D.** A zoomed-in view of the tripartite interface where the C-terminal tail of Cα simultaneously contacts both the Aα and B55α subunits from panel (B). **E.** A zoomed-in view of the interaction between the Aα and Cα subunits from panel (B). **F** and **G**. Side view (F) and bottom view (G) model of PP2A(B55)-ENSA^S67D^ complex (this study, PDB ID 9XGY, under release). **H.** A zoomed-in view of the interaction between the B55α and Cα subunits from panel (F). **I.** A zoomed-in view of the tripartite interface where the C-terminal tail of Cα simultaneously contacts both the Aα and B55α subunits from panel (G). **J.** A zoomed-in view of the interaction between the Aα and Cα subunits from panel (G). **F**-**H**. Mn^2+^ are colored magenta. **A-G**. The Aα, B55α, Cα, and ENSA^S67D^ are colored gray (90%), slate, lime green, and orange, respectively.

Importantly, ENSA^S67D^ binding induces a significant conformational rearrangement that locks the holoenzyme in a relatively extended, inhibited state. Upon ENSA^S67D^ binding, the Cα subunit shifts away from the B55α subunit and the N-terminus of Aα (Figure S11A, C); as a result, the B55α and Cα subunits interact loosely. Both the interaction between Cα Y91 and B55α D209, and that between Cα R89 and B55α N174, are disrupted (Figure 6F, H). The upper lip loop shifts away from the β1-β2 hairpin of B55α (Figure 6F, H). The Cα C-terminus-Aα-B55α interface is rearranged (Figure 6G, I), and the interaction between K416 in H11 of Aα and Cα D290 is also disrupted (Figure 6 E, G, J). This interaction rearrangement among the PP2A subunits upon ENSA^S67D^ binding also similarly occurs upon binding of the other inhibitors, recruiters, and substrates (Figure S11).

Although B55i and IER5 do not interact directly with the Cα subunit, they can induce a shift in the Cα subunit that is similar to the one induced by ENSA^S67D^ (Figure S11A-D). The C-terminus of the Aα subunit is slightly contracted upon ENSA^S67D^ and ARPP19 binding indicated by the distance between N and C terminus of Aα, whereas it is slightly extended outward upon IER5, B55i, FAM122A, Eya3, and p107 binding (Figure S11E-G). Despite these varied conformational changes in the C-terminus of Aα, Heat repeats 14-15 (“H14-15”) rotate inward upon the binding of all inhibitors, recruiters, and substrates (Figure S11E-G). Furthermore, the shifts of the Cα subunit are similar upon the binding of all these binders (Figure S11H, I).

### Conformational change of active site residues in the PP2A C subunit

Shi *et al.* suggested that the binding of B55i and Eya3 only slightly affects the conformation of key amino acids in the active site (21). Our comparison reveals that Eya3 has no significant effect on the conformation of key amino acids in the active site (Figure S12A). In contrast, B55i has a slight effect on the conformation of these amino acids, with a minor shift observed in the loops near D85 and N117 (Figure S12B). This trend of conformational change is consistent with that reported by Shi *et al* (21). ENSA^S67D^, tpARPP19, and FAM122A all interact directly with the active site of the Cα subunit, which results in the strong inhibitory effects of tpARPP19 and FAM122A on the PP2A(B55α) holoenzyme (16). While tpARPP19 and FAM122A have no effect on the conformation of key amino acids in the active site (Figure S12A), ENSA^S67D^ does have some effect (Figure S12C). p107 is a direct substrate of the PP2A(B55α) holoenzyme. The binding of p107 has a weaker effect on the conformation of key active site amino acids than B55i does (Figure S12D), and p107 exhibits a weak inhibitory effect on the PP2A(B55α) holoenzyme (22). IER5 also has a weaker effect on the conformation of these key amino acids than B55i does (Figure S12E).

Some binders (inhibitors/recruiters/substrates) have varying degrees of influence on the conformation of key amino acids in the active site, but it remains unclear how these changes influence the PP2A(B55) enzymatic activity (21). Future studies is needed to address this issue.

## Discussion

### The inhibiting model for the ENSA on PP2A(B55α) holoenzyme

Our findings provide a structural and mechanistic framework for understanding how ENSA functions as an inhibitor of PP2A(B55α). We propose a dynamic model in which the PP2A(B55α) holoenzyme transitions between two principal conformational and functional states: a “contracted” active state and an “extended” inactive state (Figure S13). During mitotic entry, pENSA acts as an inhibitor by interacting with both the B55α and Cα subunits. Although pENSA does not directly interact with the Aα subunit, its binding ultimately causes an inward rotation of HEAT repeats 14-15 (H14-15) of the Aα subunit. This, in turn, shifts the Cα subunit away from the B55α subunit and the N-terminus of Aα, causing the PP2A(B55α) holoenzyme to transition from the contracted state to the extended state. The insertion of the pS67-containing motif into the Cα catalytic site locks the PP2A(B55α) holoenzyme in this extended, inactive conformation. This inhibition is crucial for driving mitotic progression.

Although B55i (21, 26) and IER5 (23) do not interact directly with the Cα subunit, they can induce a shift in the Cα subunit that is similar to the one induced by ENSA^S67D^. This suggests that the dynamic processes by which different binders (inhibitors (16), recruiters (21–23), or substrates (22)) may trigger similar conformational changes in PP2A(B55α). This causes an inward rotation of H14-15 of the Aα subunit, ultimately leading to a positional translation of the entire Cα subunit. Simultaneously, the upper lip loop in the Cα subunit moves away from the β1-β2 hairpin of B55α, exposing the PP2A(B55α) active site. Inhibitors such as ENSA, ARPP19, and FAM122A bind to this exposed active site via a helical motif, preventing substrate binding. In contrast, recruiters like Eya3 and IER5 utilize this state to recruit PP2A(B55α) to their respective substrates, allowing the substrate’s phosphate group to bind to the exposed active site for dephosphorylation. Substrates that can directly interact with the B55α subunit (e.g., p170) can also directly dock their phosphate sites onto the exposed active site to be dephosphorylated.

### Structural plasticity of PP2A(B55α) offers avenues for novel binders discovery

Our comparative analysis reveals that the PP2A(B55α) holoenzyme exhibits remarkable plasticity in its recognition of endogenous binders (inhibitors, recruiters, or substrates), providing a rich landscape for the discovery of novel therapeutic agents. We identified five highly conserved surfaces on the B55α subunit, including two previously uncharacterized surfaces (Surfaces 4 and 5), which represent potential druggable pockets.

The known inhibitors leverage these surfaces with striking diversity. While vertebrate ENSA and ARPP19 employ a multi-site engagement strategy (Surfaces 1-3), other orthologs like *Ce*-ENSA use a different combination (Surfaces 1, 2, and 5). In contrast, FAM122A achieves potent inhibition through a highly focused interaction exclusively with Surface 2. The uniqueness of FAM122A is further underscored by its distinct SliM sequence and its fundamentally different mode of engagement with both B55α and the Cα subunit.

This diversity demonstrates that PP2A(B55α) can be allosterically modulated through multiple, distinct mechanisms. For example, Eya3 recruits the PP2A(B55α) holoenzyme to dephosphorylate Myc at pT58 (27). Given that PP2A(B55α) has numerous substrates (1), it is likely that other, yet-to-be-discovered recruiters exist.

In this study, we have newly defined a conserved surface on B55α, termed Surface 4. None of the previously reported structures of PP2A(B55α) in complex with binders (inhibitors, recruiters, or substrates) (16, 21–23) show any interaction with this surface. We therefore propose that our newly discovered Surface 4 may serve as a potential binding platform for novel PP2A(B55α) holoenzyme binders. A much-awaited PP2A(B55α) recruiter might, similar to Eya3, present the holoenzyme near potential substrates. Alternatively, such a recruiter could bind to Surface 4 at one end and to another regulator at the other, ultimately assisting the PP2A(B55α) holoenzyme in catalyzing the dephosphorylation of specific substrates. Crucially, the newly identified Surface 4, much like the well-established Surface 2 (21, 26), remains an unexploited surface that presents a prime target for the rational design of novel and specific inhibitors. Together, our findings not only suggest that additional endogenous regulators may exist but also provide a structural blueprint for developing new classes of PP2A(B55α) modulators that exploit this multifaceted interaction landscape.

### Probing the dynamics and broader context of PP2A regulation

This work lays the groundwork for several exciting avenues of future research. Phosphorylation at Ser109 within the C-terminal tail of ENSA may accelerate pS67 dephosphorylation (11), suggesting an allosteric crosstalk between the C-terminus and the catalytic core. Capturing the structure of a C-terminally truncated S109-modified ENSA complex could provide a crucial snapshot of the substrate-handoff state, further illuminating the catalytic cycle.

Furthermore, regulation of PP2A occurs within a complex network in the cell. Different inhibitors, such as FAM122A and ARPP19, can cooperate to orchestrate mitotic events (28). It will be essential to determine if similar functional coupling exists between ENSA and other regulatory proteins. Addressing these points will be key to building a comprehensive, systems-level understanding of how phosphatase activity is dynamically regulated in time and space to ensure cellular homeostasis.

## Materials and Methods

### Plasmid construction

Full-length human PP2A Aα subunit was cloned into the pGEX-4T1 vector, with an N-terminal GST tag and a TEVase cleavage site (hereafter referred to as TEV site), and expressed as previously reported (29). The full-length human PP2A B55α subunit was cloned into pFastBac vector with an N-terminal His_10_-tag and a TEV site. Full-length human PP2A Cα subunit (residues 2-309) was cloned into pDEST8 vector with an N-terminal His_8_-tag. Full-length human ENSA^WT/mutants^ were cloned into pET-28a vector with an N-terminal His_6_-tag and a TEV site. Additionally, full-length human ENSA^WT/mutants/truncations^ were cloned into pGEX-4T1 vector with an N-terminal GST tag, an N-terminal TEV site, and an additional C-terminal His_6_-tag.

### Protein expression

An optimized recombinant TEVase protease was expressed and purified as previously reported (30). Hi-5 cells, maintained in 800 ml of SF 900 II SFM (ThermoFisher), were grown to a density of 2.0 × 10^6^ cells ml^-1^. Cells were infected with 2-3% (v/v) P3 recombinant baculovirus encoding either B55α (His_10_-TEV site-B55α) or Cα (His_8_-Cα), and incubated at 27 °C for 48 h, aiming for a 10-15% cell death ratio. The Hi-5 cells were then collected by centrifugation at 800 g for 15 minutes. Cell pellets were resuspended in 50 mM Tris-HCl (pH 8.0), 500 mM NaCl, 5 mM β-mercaptoethanol (β-ME) supplemented with protease inhibitors (1× Cocktail) and stored at -80 °C. ENSA protein was expressed in *E. coli* strain ER2566. Expression was induced with 0.2 mM IPTG for 18 hours at 16 °C.

### Protein purification

Recombinant Aα purification followed a procedure similar to that previously reported (29).

For B55α purification, thawed B55α cells were resuspended in lysis buffer containing 50 mM Tris-HCl (pH 8.0), 500 mM NaCl, 5 mM β-ME, 5% glycerol, 0.1% Triton, 1× protease inhibitor cocktail, and 10× DNase. Cells were lysed by mild sonication and centrifuged twice at 17,000 rpm for 1 hour. The supernatant was filtered with a 0.8 µm syringe filter, and 30 mM imidazole was added to prevent non-specific binding. The supernatant was then incubated with Ni-NTA resin for

0.5 h at 4 °C with gentle rotation. The resin was sequentially washed with 50 mM Tris-HCl (pH 8.0), 500 mM NaCl, 5 mM β-ME, 50 mM imidazole, followed by 50 mM Tris-HCl (pH 8.0), 500 mM NaCl, 5 mM β-ME, and 100 mM imidazole. Bound proteins were eluted using 50 mM Tris-HCl (pH 8.0), 200 mM NaCl, 5 mM β-ME, and 300 mM imidazole, and 2 mM DTT was immediately added to prevent protein oxidation. The NaCl concentration of the eluted sample was adjusted to 50 mM and applied to a Q(8.0) anion exchange column pre-equilibrated with 50 mM Tris-HCl (pH 8.0), 50 mM NaCl, 2 mM DTT. Bound proteins were concentrated by one-step elution (Q(8.0) (5-100%B, 0 mL)). The eluted fraction was further purified by size-exclusion chromatography (SEC) using a Superdex 200 increase 10/300 column (Cytiva) pre-equilibrated with 20 mM HEPES (pH 8.0), 100 mM NaCl, 2 mM DTT. The purified B55α was used for complex assembly and other experiments.

For Cα purification, thawed Cα cells were resuspended in lysis buffer containing 50 mM Tris-HCl (pH 8.0), 500 mM NaCl, 5 mM β-ME, 1× protease inhibitor cocktail, and 10× DNase. Cells were lysed by mild sonication and centrifuged at 17,000 rpm for 1 hour. The supernatant was filtered with a 0.8 µm syringe filter, and 20 mM imidazole was added to prevent non-specific binding. The supernatant was then incubated with Ni-NTA resin for 0.5 h at 4 °C with gentle rotation. The resin was washed with 50 mM Tris-HCl (pH 8.0), 500 mM NaCl, 5 mM β-ME, and 50 mM imidazole. Bound proteins were eluted with 50 mM Tris-HCl (pH 8.0), 200 mM NaCl, 5 mM β-ME, and 300 mM imidazole, and 2 mM DTT was immediately added to prevent protein oxidation. The NaCl concentration of the eluted sample was adjusted to 25 mM and applied to a Q(8.0) anion exchange column pre-equilibrated with 50 mM Tris-HCl (pH 8.0), 25 mM NaCl, 2 mM DTT. Bound proteins were concentrated by one-step elution (Q(8.0) (2.5-100%B, 0 mL)). The eluted fraction was further purified by SEC using a Superdex 200 increase 10/300 column (Cytiva) pre-equilibrated with 20 mM HEPES (pH 8.0), 100 mM NaCl, 2 mM DTT. The purified Cα was used for complex assembly and other experiments.

For ENSA protein purification, thawed ENSA cell pellets were resuspended in lysis buffer containing 50 mM Tris-HCl (pH 8.0), 500 mM NaCl, 5 mM β-ME, 1× PMSF, and 1× DNase. Cells were lysed by sonication and centrifuged at 17,000 rpm for 1 h. The supernatant was filtered with a 0.8 µm syringe filter, and 20 mM imidazole was added to prevent non-specific binding. The supernatant was then incubated with Ni-NTA resin for 0.5 h at 4 °C with gentle rotation. The resin was washed with 50 mM Tris-HCl (pH 8.0), 500 mM NaCl, 5 mM β-ME, and 50 mM imidazole. Bound His_6_-TEV site-ENSA^WT/mutants^ and GST-ENSA^WT/mutants/truncations^-His_6_ proteins were eluted with 50 mM Tris-HCl (pH 8.0), 200 mM NaCl, 5 mM β-ME, and 300 mM imidazole, and 2 mM DTT was added to prevent protein oxidation. The eluted fraction was further purified by SEC using a Superdex 200 increase 10/300 column (Cytiva) pre-equilibrated with 20 mM HEPES (pH 8.0), 100 mM NaCl, 2 mM DTT.

For Microscale thermophoresis (MST) and Isothermal titration calorimetry (ITC) assays, His_6_-TEV site-ENSA^WT/mutants^ proteins were incubated with TEVase protease (1:30 molar ratio) at room temperature for 1 h. The mixture was then reapplied to Ni-NTA resin for 0.5 h at 4 °C to remove cleaved His_6_-TEV site fragments. The tag-free ENSA^WT/mutants^ (with an N-terminal G amino acid residue, the same position is S, as previously reported (13)) in the flow-through were further purified by SEC using a Superdex 200 increase column pre-equilibrated with 20 mM HEPES (pH 8.0), 100 mM NaCl, 2 mM DTT.

### Protein complex assembly and purification

For cryo-EM analysis of the PP2A-ENSA^S67D^ complex, purified PP2A Aα, B55α, and Cα subunits were initially incubated in a 1:1.3:1.1 molar ratio for 2 h at 4 °C with gentle rotation. Subsequently, His_6_-TEV site-ENSA^S67D^ protein was added to a 20-fold molar excess (relative to Aα) and the mixture was further incubated for 4 h at 4 °C with gentle rotation. The sample was then applied to a Superdex 200 increase column (Cytiva) pre-equilibrated with 20 mM HEPES (pH 8.0), 100 mM NaCl, and 2 mM DTT. An additional 5-fold molar excess of His_6_-TEV site-ENSA^S67D^ protein was added to the purified complex fractions to stabilize the PP2A-ENSA^S67D^ complex.

To assemble the apo PP2A(B55) holoenzyme complex, Aα, B55α, and Cα subunits were incubated in a 1:1.7:1.2 molar ratio for 0.5 h at 4 °C with gentle rotation. The sample was then applied to a Superdex 200 increase column (Cytiva) pre-equilibrated with 20 mM HEPES (pH 8.0), 100 mM NaCl, and 2 mM DTT.

### Cryo-EM sample preparation and data acquisition

For cryo-EM analysis, a 3.0 µL drop of the 1.6 mg/mL PP2A-ENSA^S67D^ complex (with a 5-fold molar excess of ENSA^S67D^) was loaded onto a glow-discharged holey carbon grid (Quantifoil, Cu R1.2/1.3, 300 mesh), which had been treated with a Gatan Plasma System (H_2_/O_2_ for 40 s). The grids were blotted for 5.0 s with a force of -1.0 and then plunge-frozen into liquid ethane using a ThermoFisher Vitrobot Mark IV (humidity 100%, 10 °C, blotting paper TED PELLA 595).

Data collection was automatically performed using SerialEM (31). Movies were acquired on a Titan Krios G3 microscope (ThermoFisher) operating at 300 kV. The microscope was equipped with a K3 Summit direct electron detector (Gatan) and a Quantum energy filter (Gatan), with a slit width of 20 eV. Movies were recorded in super-resolution mode at a nominal magnification of ×105,000, corresponding to a pixel size of 0.832 Å. The defocus range was set from -1.0 to -2.0 µm. All movies were exposed at a total dose of 60 e^-^/Å² over 40 frames.

### Cryo-EM data processing

For the dataset of the PP2A-ENSA^S67D^, 5,579 raw movies were motion-corrected and dose-weighted using Relion 4.0 (32), and their contrast transfer functions were estimated by cryoSPARC patch CTF estimation (33). After curating exposures, 4,110 selected images were conducted the blob, template and topaz (34) picker jobs. Following 2 round of 2D classifications, 828,786 particles were selected and performed ab-Initio reconstruction in four classes. These four classes then served as 3D volume templates for heterogeneous refinement with all selected particles. One good 3D class including 543,811 particles was used to perform non-uniform refinement (NU-refinement) and CTF refinement, yielding a resolution of 2.85 Å with poor density of the ENSA^S67D^. To enhance the quality of the ENSA^S67D^ density, we manually created a mask which includes the ENSA^S67D^ and part of the PP2A density. Subsequently, we performed no-alignment 3D classification with the manually-created mask to locally classify, selecting a class that contained the most complete ENSA density, comprising 227,045 particles. We then utilized NU-refinement to reconstruct these particles, resulting in a map with an overall resolution of 3.03 Å. In this map, the density of the ENSA region was significantly improved (Figure S3).

The reported resolutions above are based on the gold-standard Fourier shell correlation (FSC) 0.143 criterion. All the visualization and evaluation of 3D density maps were performed with UCSF Chimera (35) and ChimeraX (36). The final sharpened map was generated using EMReady (24) to preserve optimal complete density, and verified for correct handedness in UCSF Chimera prior to subsequent model building and analysis.

### Model building and refinement

An AlphaFold3-predicted model (25) of the PP2A-ENSA^S67D^ complex was fitted into the cryo-EM map with Phenix (37). The initial model was manually adjusted in Coot (38) and subsequently refined in Phenix (37) for several rounds. The protein-protein interfaces were analyzed with PDBePISA. All structure representations were generated with ChimeraX (36) and PyMOL 3.0 for macOS (http://www.pymol.org).

### GST pull-down

Approximately 0.92 μg GST, 1.47 μg GST-ENSA^WT/S67D^-His_6_, 11.48 μg Aα, 9.15 μg B55α and 6.54 μg Cα were used in the pull down assay, respectively. The molar ratio of GST, GST-ENSA^WT/S67D^-His_6_, PP2A(B55α) Aα subunit, B55α subunit, AC core enzyme and holoenzyme was 1: 1: 5: 5: 5: 5.

PP2A related proteins were mixed in 50 µl of pull-down buffer containing 20 mM HEPES (pH 8.0), 150 mM NaCl, 2 mM DTT, 0.05% (v/v) tween 20 and incubated at 4°C for 1 h. Subsequently, GST and GST-ENSA^WT/S67D^-His_6_ was added and the mixture was incubated for an additional 2 h at 4°C. Glutathione resin (20 μL) and 130 μL of pull-down buffer were then added, followed by a further 2 h incubation at 4 °C. The resin was washed three times with 180 μL of pull-down buffer. Bound proteins were analysed by sodium dodecyl sulfate polyacrylamide gel electrophoresis (SDS-PAGE) and stained with Coomassie blue.

### Microscale thermophoresis assays

The MST assay were performed as previously reported (13). The binding affinities between tag-free ENSA^WT/S67D^ and PP2A Aα subunit were measured using a Monolith NT.115 (NanoTemper Technologies). The Aα protein was labelled with Protein Labeling Kit RED-NHS 2nd Generation (Cat no. MO-L011, NanoTemper Technologies) according to the operation manual. The labelled Aα protein, at a concentration of 20 nM, was incubated with the unlabelled tag-free ENSA^WT/S67D^ at 14 different concentrations (ranging from 1.8 mM to 0.2 µM) in MST buffer (50 mM NaH_2_PO_4_ (pH 6.8), 100 mM KCl, 1 mM DTT, 0.05% Tween-20, 0.5 mg/ml BSA) at room temperature for 30 min. The protein mixtures were then loaded into capillaries (Cat no. MO-K022, NanoTemper Technologies) and measured at 25 °C using medium MST power and 20% Excitation power. Each assay was repeated three times. Dissociation constant (*K*_D_) values were calculated using the GraphPad Prism 10.2.3 for macOS.

### Isothermal titration calorimetry assays

ITC was performed using a MicroCal PEAQ-ITC (Malvern) at 25 °C. Proteins were buffer exchanged into 20 mM HEPES (pH 8.0), 100 mM NaCl, 4 mM β-ME via SEC on a Superdex 75 increase 10/300 column (Cytiva). Approximately 1000 µM tag-free ENSA^S67D^ (titrant) was injected into 50 µM PP2A Aα subunit (titrand) with 18 injections of 2 µL each, at 2.5-min intervals. Similarly, 1000 µM tag-free ENSA^WT^ was injected into 100 µM Aα subunit under the same titration conditions. Data were fitted to a one-site model using the MicroCal PEAQ-ITC analysis software 1.4 (Malvern).

### Bio-layer interferometry (BLI) assays

BLI was performed using an Octet RED96 instrument (FortéBio) at 25 °C. All experiments utilized BLI buffer containing 20 mM HEPES (pH 8.0), 100 mM NaCl, 2 mM DTT, and 10 mg/mL BSA. Single referencing was applied by subtracting signals from a buffer-only control. Double referencing involved subtracting signals from both reference sensors and buffer-only controls. GST-ENSA^WT/mutants/truncations^-His_6_ were mobilized on anti-GST biosensors (Sartorius).

The interaction between GST-ENSA^WT/S67D^-His_6_ and PP2A-related proteins (PP2A Aα subunit, Cα subunit and AC core enzyme) were measured using a double referencing assay. The concentration gradients of the PP2A-related proteins are indicated in the figures. Data were presented using the GraphPad Prism 10.2.3 for macOS.

For binding affinities between GST-ENSA^WT/S67D^-His_6_ and PP2A-related proteins (PP2A B55α subunit and holoenzyme), a single referencing assay was employed. The concentration gradients of PP2A-related proteins are also indicated in the figures. *K*_D_ values were calculated using FortéBio data analysis software 10.0. Data were presented using the GraphPad.

To analyze the interface arcletecture of ENSA and PP2A(B55) holoenzyme, a single referencing assay was performed with 3 µM PP2A. Data were presented using the GraphPad.

### Conserved amino acid analysis and sequence alignment of proteins

Conserved amino acid analysis of Aα and B55α subunits across 150 organisms was performed using the ConSurf server (39) with sequence-only input. The conservation scores from ConSurf were visualized on the 3D structure of the protein using PyMOL 3.0 for macOS.

The sequence alignment of proteins were performed using T-Coffee (40, 41) and visualized with Jalview 2.11.4.1 (42) for macOS.

### Data, materials, and software availability

Cryo-EM data for the PP2A(B55)-ENSA^S67D^ complex have been deposited in the Protein Data Bank (PDB) (https://www.rcsb.org) under accession code 9XGY (43) and the Electron Microscopy Data Bank (EMDB) (https://www.ebi.ac.uk/emdb/) under accession code EMD-66862 (44). All other data are included inthe article and/or *SI Appendix*.

## Acknowledgments

We thank Qianqian Sun and Li Wang at the Bio-Electron Microscopy Facility of ShanghaiTech University for technical assistance with cryo-EM data collection. We thank the Cryo-Electron Microscopy Center of the Zhongshan Institute for Drug Discovery (ZIDD) for providing support and assistance in Cryo-EM data analysis. We thank the Molecular and Cell Biology Core Facility (MCBCF) at the School of Life Science and Technology, ShanghaiTech University for providing technical support of MST assays. We thank Yi Zhang from the Discovery Technology Platform at the Shanghai Institute for Advanced Immunochemical Studies (SIAIS), ShanghaiTech University for providing technical support of Octet/BLI assays. We thank Xiuxia Gao from the Analytical Chemistry Platform at the Shanghai Institute for Advanced Immunochemical Studies (SIAIS), ShanghaiTech University, for assistance with ITC analysis. We thank the High-Performance Computing (HPC) Platform of ShanghaiTech University for its support with computational resources. This work was supported by the National Key R&D Program of China Grant (2024YFA0916900 to W.X.), and the Chinese Academy of Sciences Pilot Strategic Science and Technology Project B Grant (XDB37030302 to W.X.), as well as a startup fund from ShanghaiTech University (to W.X.). Additional support was provided by the Shanghai Frontiers Science Center for Biomacromolecules and Precision Medicine at ShanghaiTech University.

## Author contributions

G.T., Z.W. and W.X. designed research. G.T. and H.L. purified the protein complexes. G.T. prepared the cryo-EM samples, and collected cryo-EM data. X.Z. and G.T. processed cryo-EM data. G.T., X.Z. and Z.W. refined the density maps and built the atomic models. G.T. conducted the biochemical experiments. G.T. and G.S. processed computational analysis. L.X., Y.Y., Q.M., Y.G., L.S. and G.X. contributed to analysis and discussion. G.T., Z.W. and W.X. wrote the paper, and all authors revised the manuscript.

