## Supplementary information for "Structural and biochemical studies of human PP2A(B55) holoenzyme and ENSA protein complex"

**This PDF file includes:**

Figures S1 to S13
Tables S1 to S2

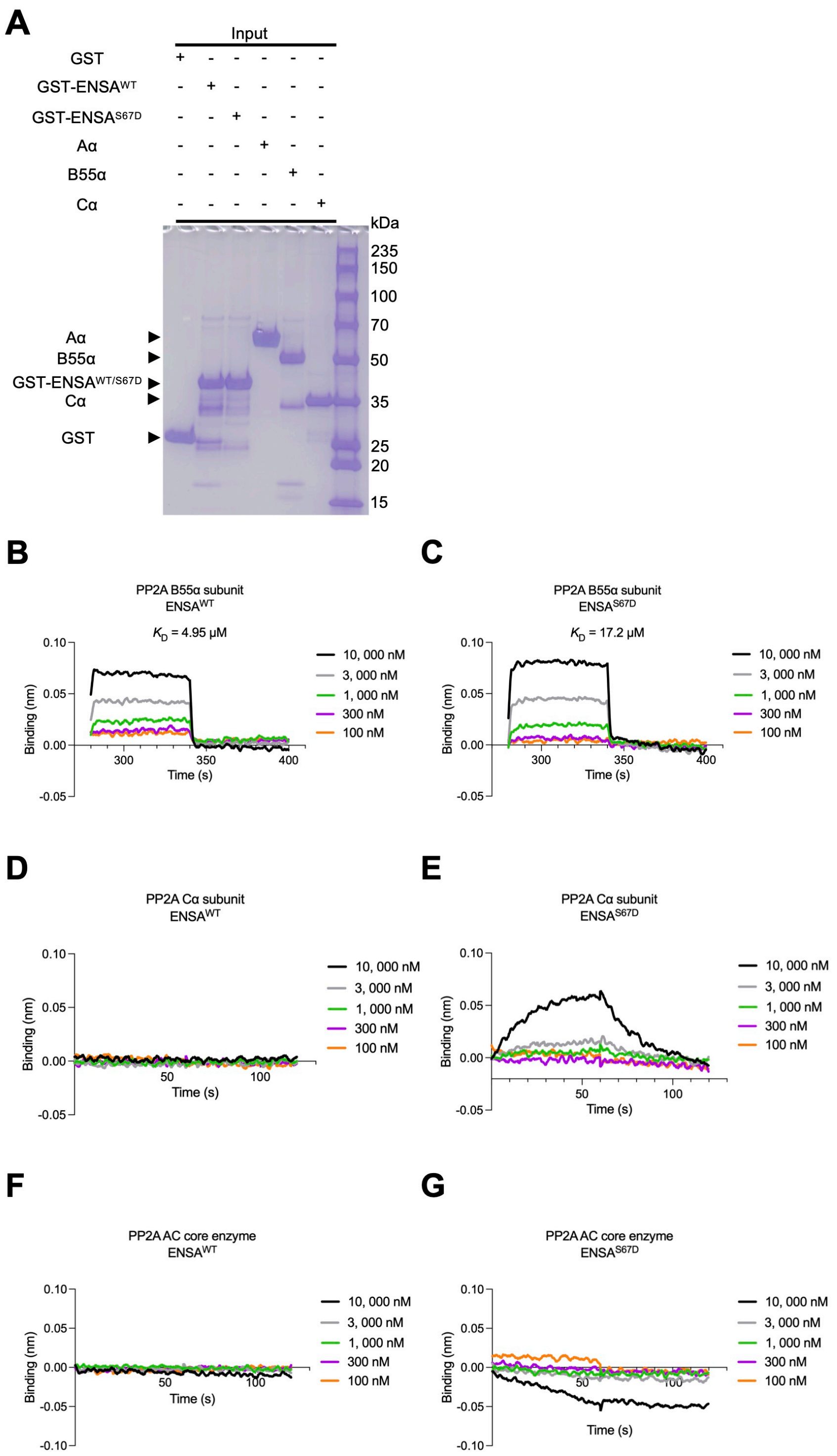


**Figure S1. Assessment of potential interactions of ENSA$^{WT/S67D}$ to PP2A(B55) AC core enzyme and subunits, using BLI.**

**A**, Inputs for GST pull-down assays in Figure 1A. Loading amounts of GST and GST-ENSA$^{WT/S67D}$-His$_6$ are the same as in the pull-down experiments, whereas the amounts of PP2A(B55α) Aα subunit, B55α subunit, AC core enzyme, and holoenzyme are 1/5 of the input used for the pull-down assays.

**B**-**G**, Biolayer interferometry (BLI) measuring the binding of wild-type ENSA (ENSA$^{WT}$) or the phosphomimetic mutant ENSA$^{S67D}$ to various components of the PP2A(B55α)

holoenzyme.
**B** and **C**. Interaction of the isolated B55α subunit with $ENSA^{WT}$ (B) and $ENSA^{S67D}$ (C).
**D** and **E**. Interaction of the isolated Cα subunit with $ENSA^{WT}$ (D) and $ENSA^{S67D}$ (E).
**F** and **G**. Interaction of the A-C core enzyme with $ENSA^{WT}$ (F) and $ENSA^{S67D}$ (G). Dissociation constants ($K_D$) derived from these experiments are summarized in Table S1. Details of data collection and processing are provided in the Methods.

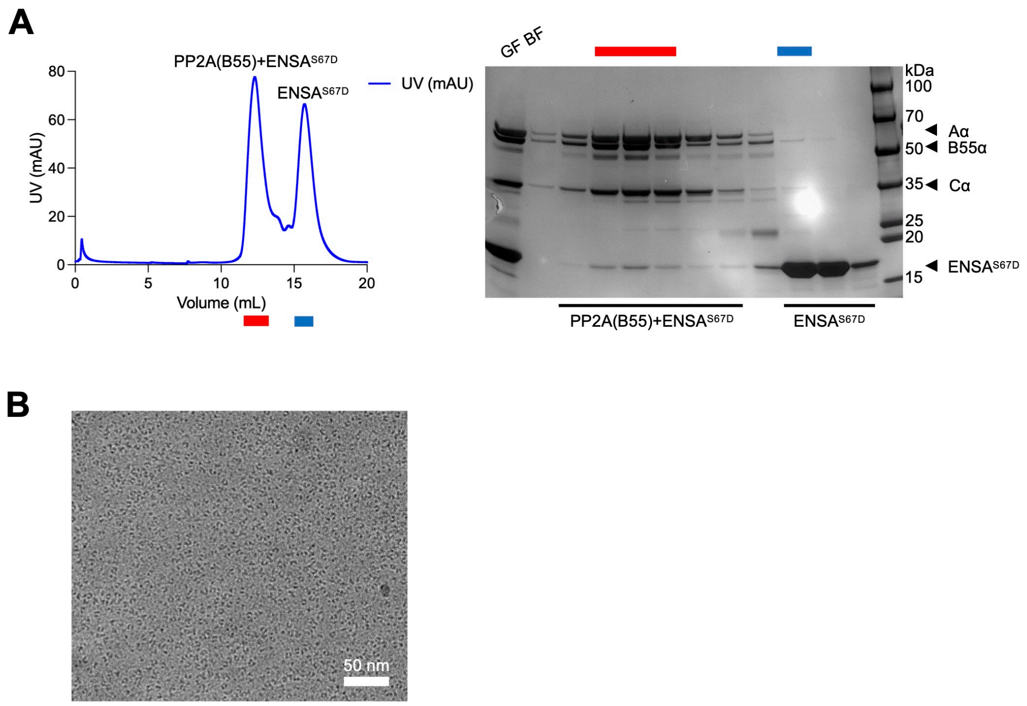


**Figure S2. Preparation and characterization of PP2A(B55)-ENSA$^{S67D}$ complexes.**

**A**. Final purification step of the PP2A(B55)-ENSA$^{S67D}$ complex by size-exclusion chromatography (SEC). The chromatogram shows a single, monodisperse peak corresponding to the fully assembled quaternary complex. The sodium dodecyl sulfate polyacrylamide gel electrophoresis (SDS-PAGE) gel confirms the presence of all four components (Aα, B55α, Cα, and ENSA$^{S67D}$) in the peak fractions (indicated by a red bar). GF BF refers to the sample loaded onto the column.

**B**. A representative cryo-EM micrograph of the vitrified complex, showing a high density of well-distributed, monodisperse particles suitable for single-particle analysis.

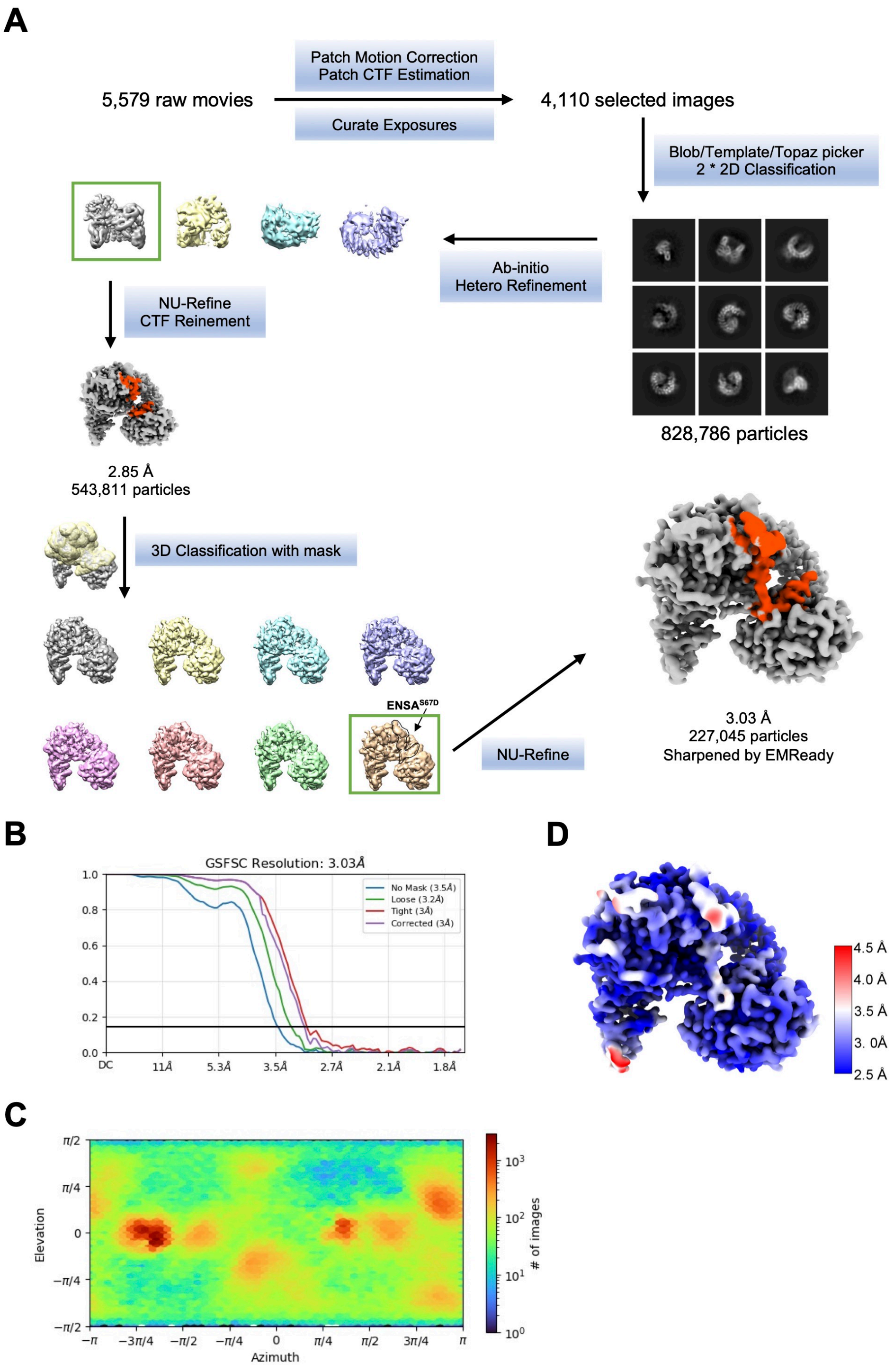


**Figure S3. The cryo-EM data processing.**

**A**, Flow chart for the processing of the cryo-EM data. Data processing details was shown in Methods.

**B**, Gold-standard fourier correlation curves of the 3D reconstructions.

**C**, Posterior precision directional distributions of all particles used in the final 3D reconstruction reported by cryoSPARC.

**D**, The color-coded local resolution map.

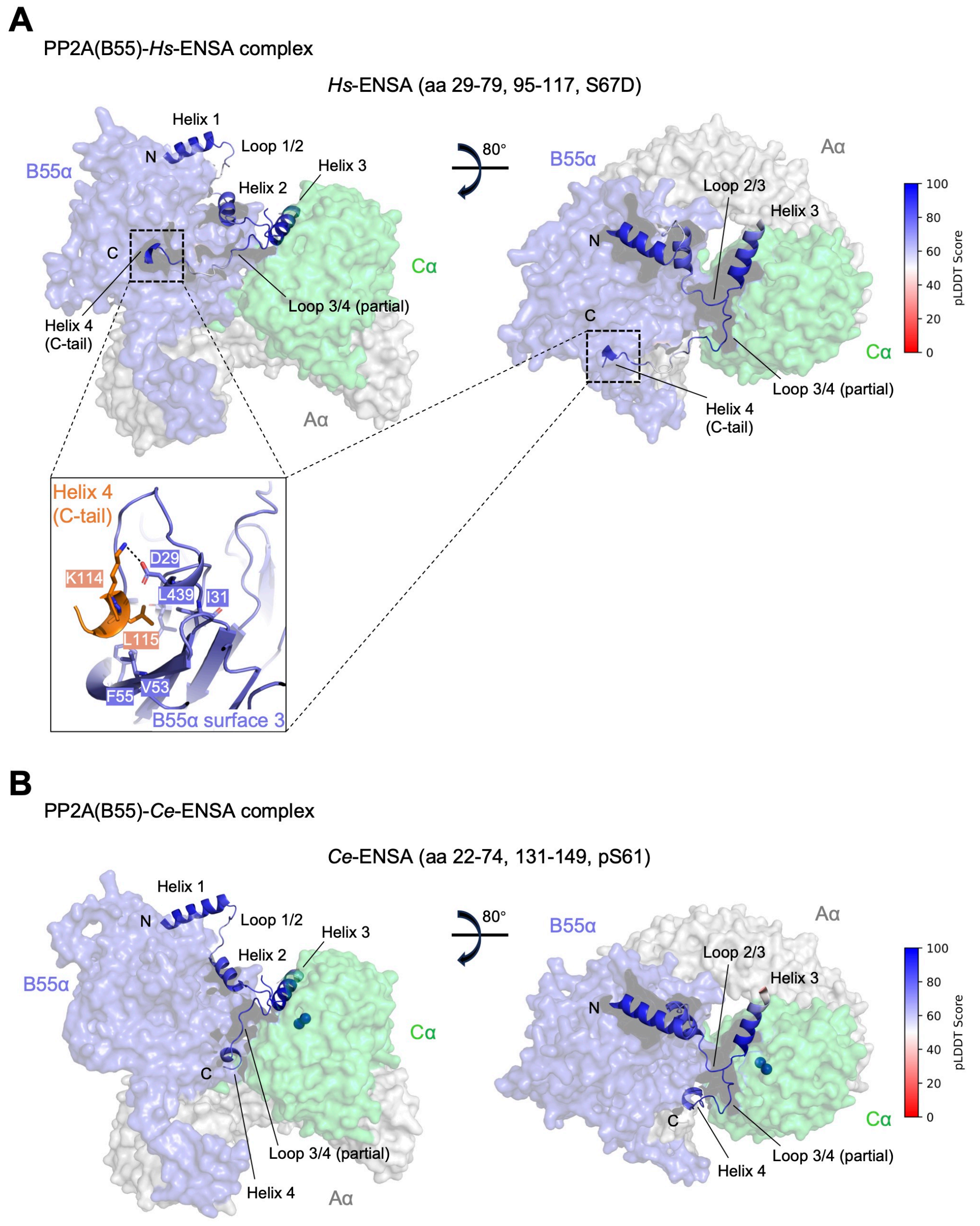


**Figure S4. Structure prediction of the potential helix-4 region of *Hs*-ENSA and *Ce*-ENSA in complex with PP2A(B55) holoenzyme.**

In both panels, the PP2A(B55) holoenzyme is shown in a semi-transparent (60%) surface representation, with subunits colored as follows: Aα (gray), B55α (slate), and Cα (lime green). *Hs*, *Homo sapiens*; *Ce*, *Caenorhabditis elegans*. *Hs*-ENSA and *Ce*-ENSA are shown in cartoons.

**A**, The AlphaFold3-predicted structure of the PP2A(B55)-*Hs*-ENSA complex.

**B**, The AlphaFold3-predicted structure of the PP2A(B55)-*Ce*-ENSA complex.

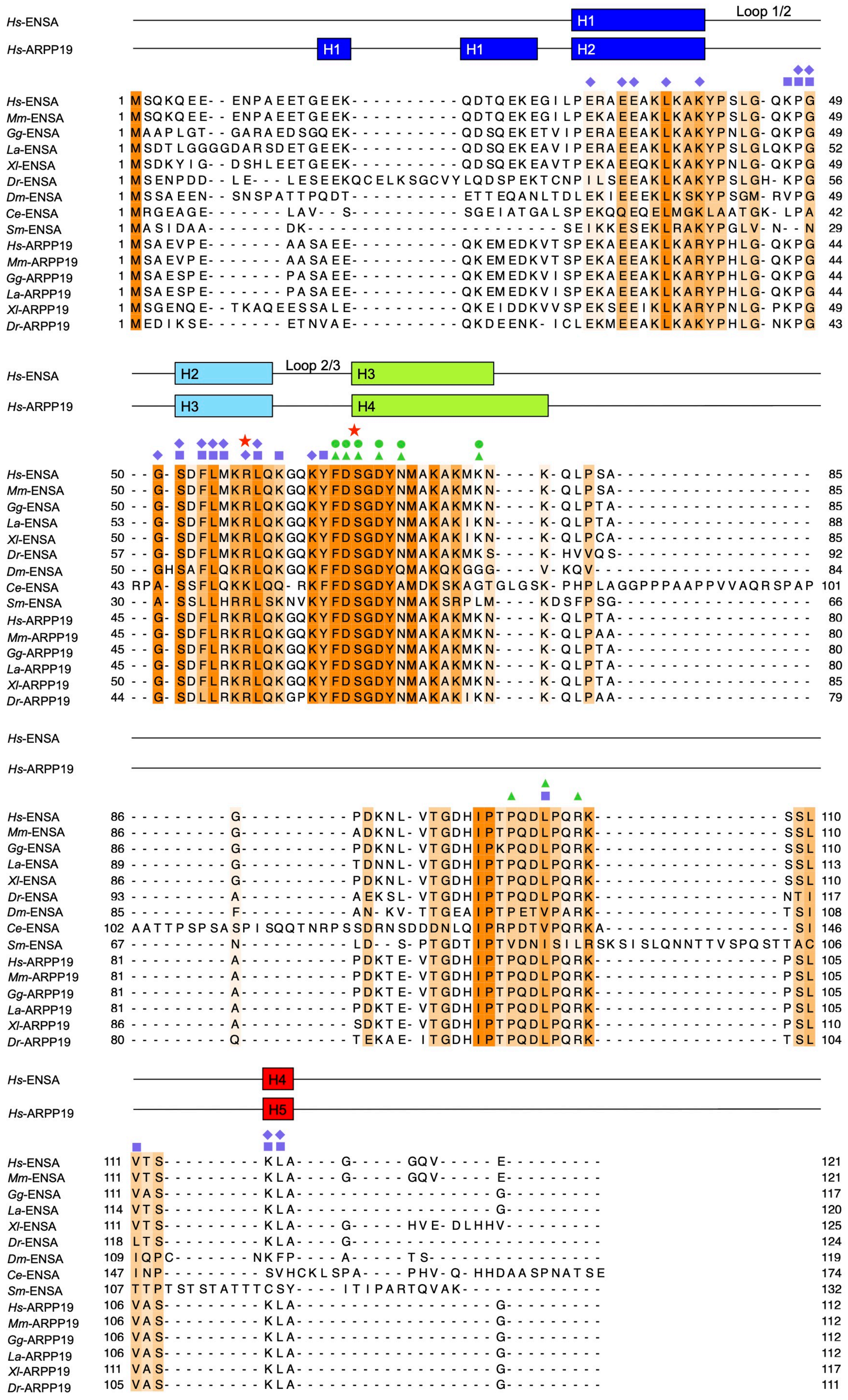


**Figure S5. The sequence alignment of ENSA and ARPP19 orthologs.**

Conserved residues between ENSA and ARPP19 are highlighted in dark orange, with the intensity corresponding to the level of conservation. Residues in *Hs*-ENSA that interact with PP2A B55α and Cα subunits are marked by medium slate blue diamonds and lime green dots, respectively. Similarly, residues in *Hs*-ARPP19 that interact with B55α and Cα subunits are marked by medium slate blue squares and lime green triangles, respectively. Key residues critical for interaction with either the B55α or Cα subunit are indicated by red stars. Organisms abbreviations are as follows: *Hs*, *Homo sapiens*; *Mm*, *Mus musculus*; *Gg*, *Gallus gallus*; *La*, *Lacerta agilis*; *Xl*, *Xenopus laevis*; *Dr*, *Danio rerio*; *Dm*, *Drosophila melanogaster*; *Ce*, *Caenorhabditis elegans*; *Sm*, *Schistosoma mansoni*.

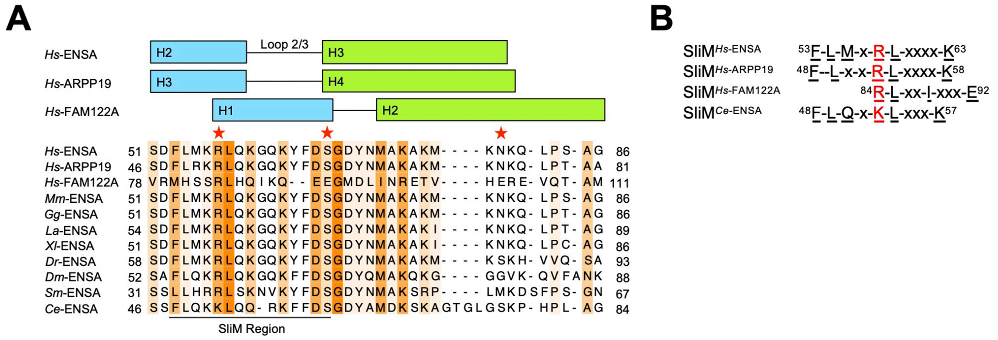


**Figure S6. The SliM region and the Cα-interacting elements of PP2A(B55) inhibitors.**

Organisms abbreviations are same to Figure S5**.**

**A**. Sequence alignment of ENSA, ARPP19, and FAM122A, focusing on the regions containing the SliM region and the Cα-interacting elements. Conserved residues are highlighted in dark orange, with the intensity corresponding to the level of conservation. The key residues that interact with B55α and Cα are marked with red stars.

**B**. A focused view of the SliM sequences from the alignment in panel (A).

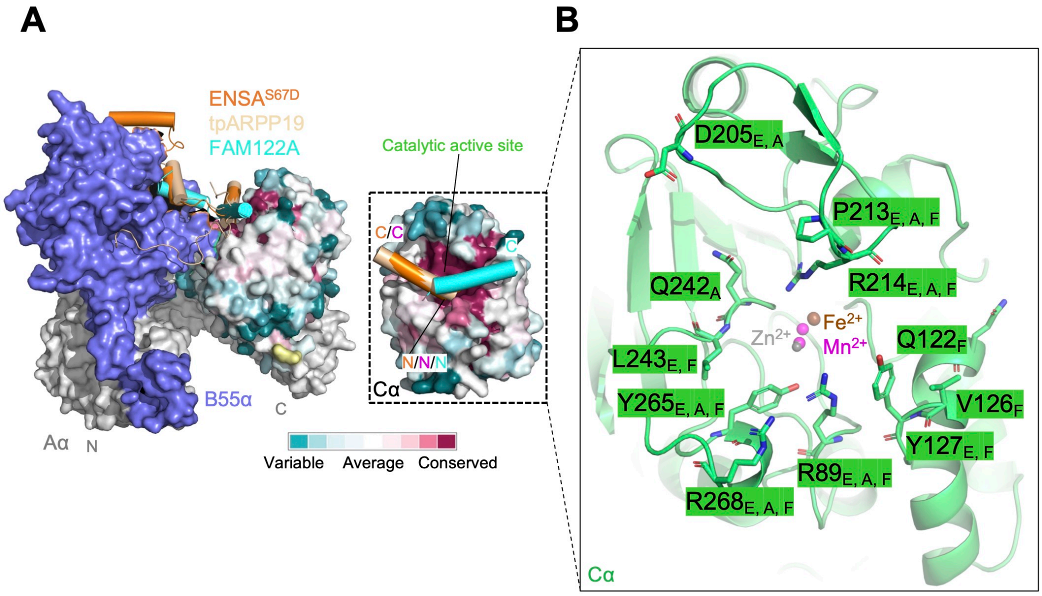


**Figure S7. Analysis of conserved PP2A-Cα subunit surface residues.**

**A**. The conserved amino acid analysis of Cα subunit across 150 organisms is generated using the ConSurf. The Cα subunit (residues 2-289) is showed as surface, while ENSA, ARPP19 and FAM122A are shown as cartoons.

**B**. Base on PP2A(B55)-ENSA$^{S67D}$ (this study), PP2A(B55)-tpARPP19 and PP2A(B55)-FAM122A, the highly conserved functional and structural residues are show as sticks. The R89$_{E, A, F}$ of Cα interacts with ENSA, ARPP19 and FAM122A. $Fe^{2+}$, $Mn^{2+}$, and $Zn^{2+}$ are shown as spheres scaled to 0.3 and colored brown, magenta, and gray (50%), respectively.

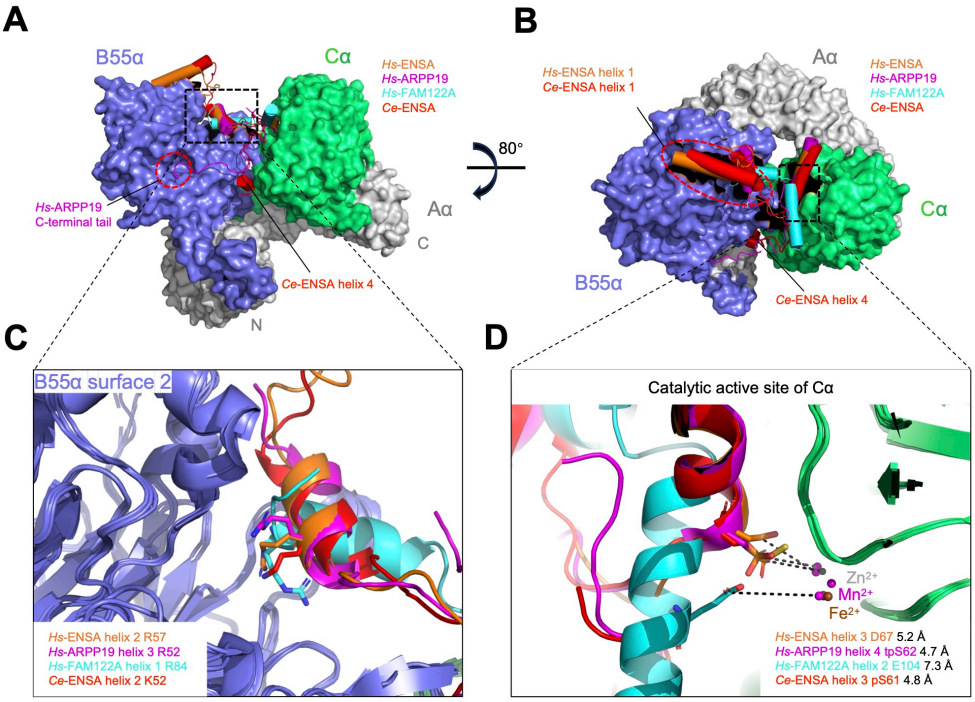


**Figure S8. Comparison of PP2A(B55) inhibitor complexes.**

**A-D**. Organisms abbreviations are as follows: *Hs*, *Homo sapiens*; *Ce*, *Caenorhabditis elegans*. **A-D**. *Hs*-ENSA, *Hs*-ARPP19, *Hs*-FAM122A, and *Ce*-ENSA are colored orange, magenta, cyan, and red, respectively.

**A** and **B**. PP2A(B55)-ENSA$^{S67D}$ cryo-EM structure (this study, PDB ID 9XGY, under release), PP2A(B55)-FAM122A (PDB ID 8SO0), *Hs* PP2A(B55)-tpARPP19 (PDB ID 8TTB), and AlphaFold3 predicted *Ce* PP2A(B55)-ENSA$^{pS61}$ are superimposed to B55α and Cα (residues 2-289) of PP2A(B55)-ENSA$^{S67D}$ structure (this study), respectively. Aα, B55α, and Cα are shown as surfaces. *Hs*-ENSA, *Hs*-ARPP19, *Hs*-FAM122A, and *Ce*-ENSA are shown as cylindrical helices.

**C**. A zoomed-in view of the interaction between the SliM sequences and Surface 2 on B55α, from panel (A). B55α, ENSA, ARPP19, and FAM122A are shown as cartoons. Conserved R and K residues are shown as sticks.

**D**. A zoomed-in view of the Cα catalytic active site, from panel (B). Cα, ENSA, ARPP19, and FAM122A are shown as cartoons. The D, S, and E residues are shown as sticks, and distance between the residues and mental iron are indicated, respectively. $Fe^{2+}$, $Mn^{2+}$, and $Zn^{2+}$ are shown as spheres scaled to 0.2 and colored brown, magenta, and gray (50%), respectively.

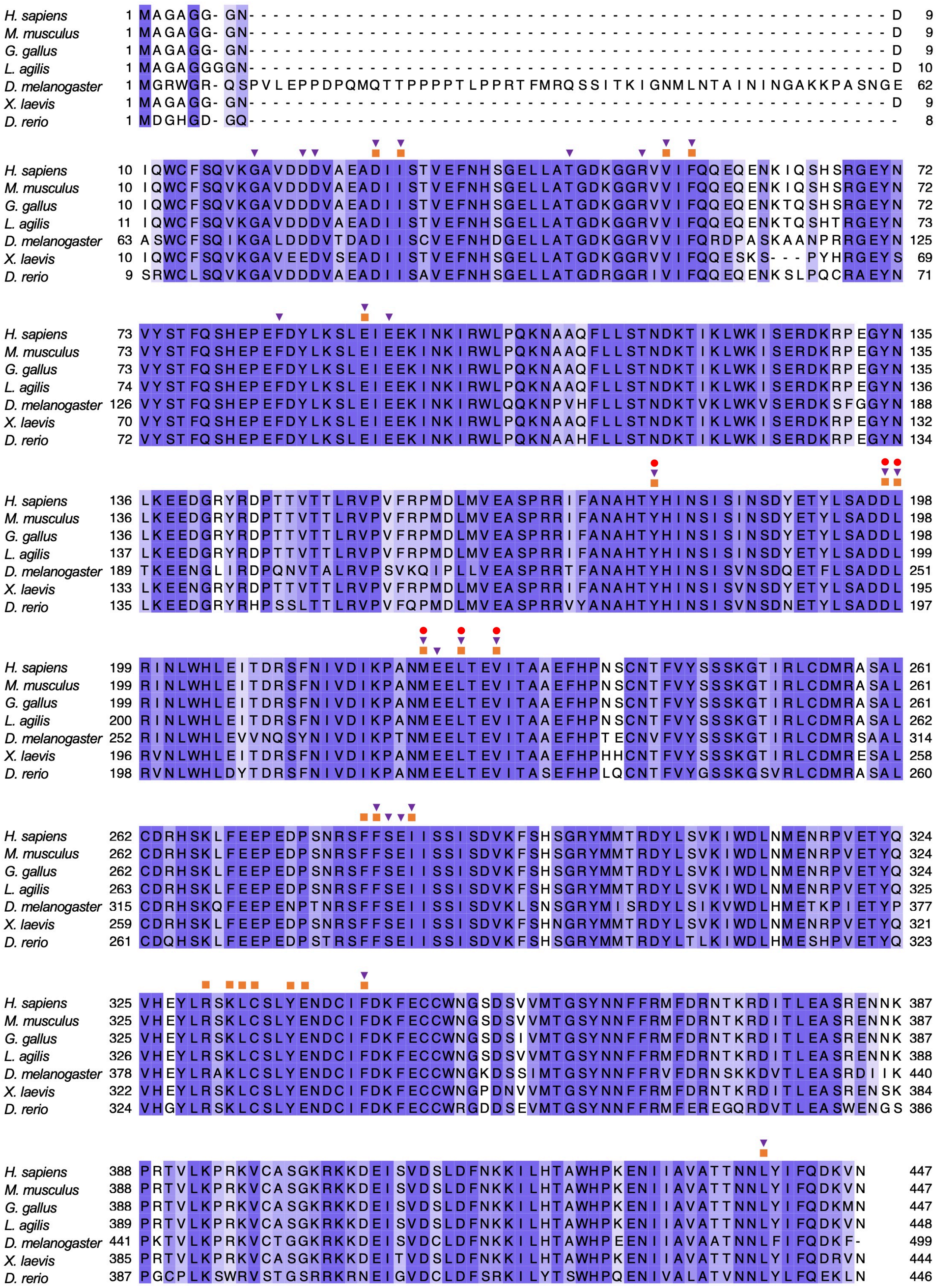

**Figure S9. The sequence alignment of B55α orthologs.**

Conserved residues in B55α is highlighted in medium slate blue, with the intensity corresponding to the level of conservation. Residues in B55α that interact with ENSA, ARPP19 and FAM122A proteins are marked by dark orange squares, medium slate blue inverted triangles, and red dots, respectively. Organisms abbreviations are as follows: *H. sapiens*, *Homo sapiens*; *M. musculus*, *Mus musculus*; *G. gallus*, *Gallus gallus*; *L. agilis*, *Lacerta agilis*; *X. laevis*, *Xenopus laevis*; *D. rerio*, *Danio rerio*.

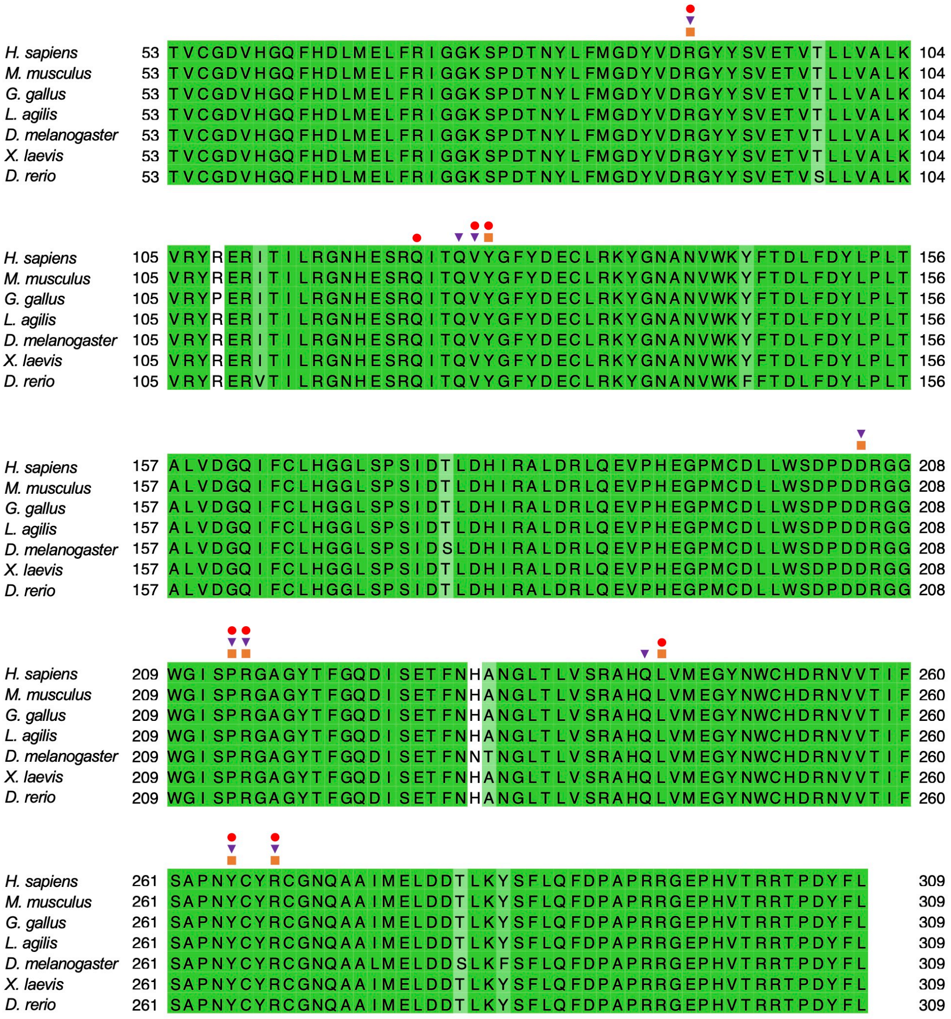


**Figure S10. The sequence alignment of Cα orthologs.**

Conserved residues in Cα is highlighted in lime green, with the intensity corresponding to the level of conservation. Residues in Cα that interact with ENSA, ARPP19 and FAM122A proteins are marked by dark orange squares, medium slate blue inverted triangles, and red dots, respectively. Organisms abbreviations are as follows: *H. sapiens*, *Homo sapiens*; *M. musculus*, *Mus musculus*; *G. gallus*, *Gallus gallus*; *L. agilis*, *Lacerta agilis*; *X. laevis*, *Xenopus laevis*; *D. rerio*, *Danio rerio*.

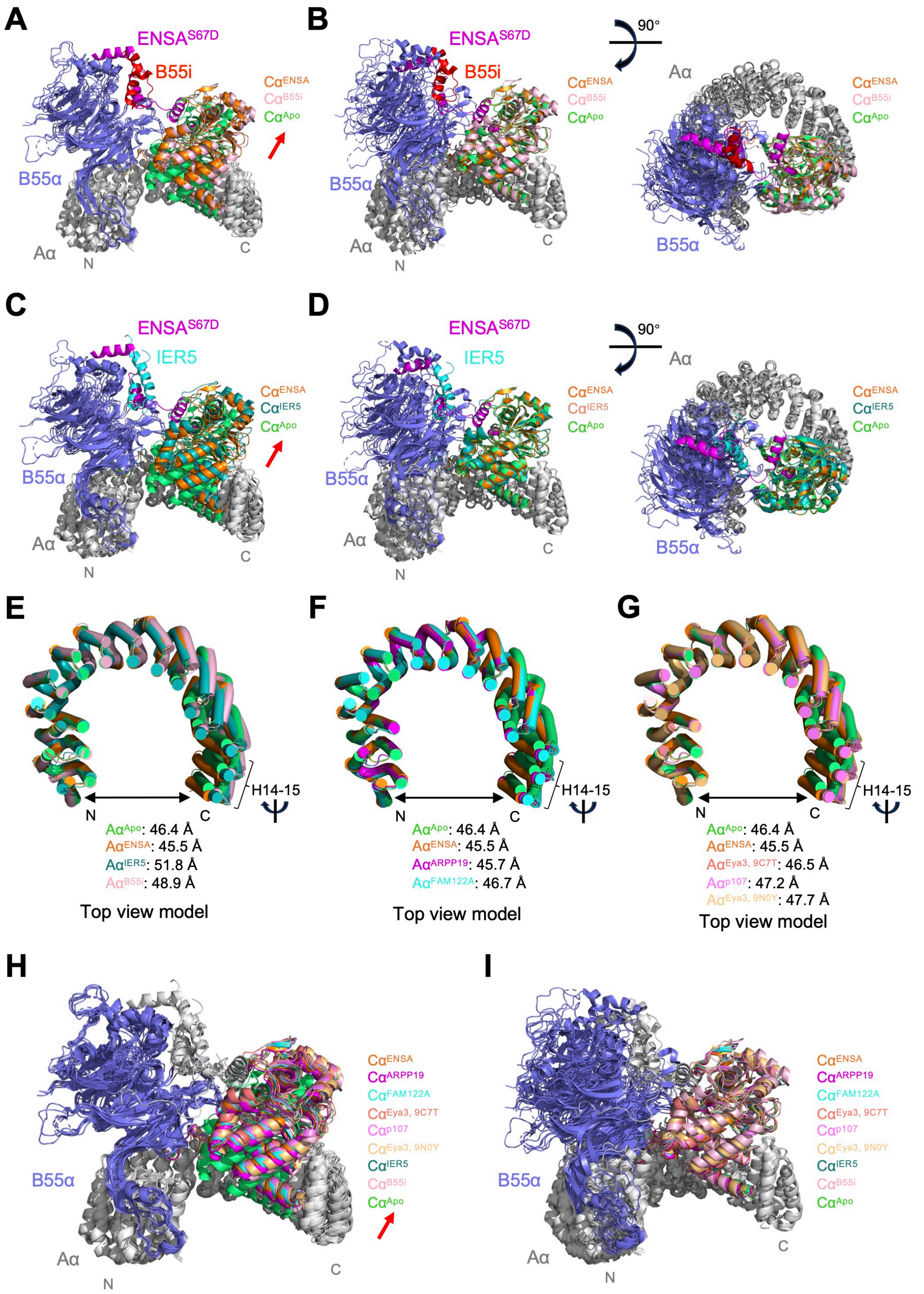


**Figure S11. The conformational changes of PP2A upon binders binding**
**A**, **C, E-G** and **H**. PP2A(B55) with differ binders are superimposed to Aa (residues 10-314) (Heat repeats 1-8) of apo PP2A(B55) (PDB ID 9MZW), respectively. **B**, **D** and **I**. PP2A(B55) with differ binders are superimposed to Aa (residues 513-589) (Heat repeats 14-15) of apo PP2A(B55) (PDB ID 9MZW), respectively. **A**-**D**. ENSA (ENSA$^{S67D}$, this study) is colored magenta. B55i (panel A and B) and IER5 (panel C and D) are colored red and cyan, respectively. **A-I**. Aα/Cα$^{ENSA}$, Aα/Cα$^{ARPP19}$, Aα/Cα$^{FAM122A}$, Aα/Cα$^{Eya3,\ 9C7T}$,

Aα/Cα$^{p107}$, Aα/Cα$^{Eya3,\ 9N0Y}$, Aα/Cα$^{IER5}$, Aα/Cα$^{B55i}$, and Aα/Cα$^{Apo}$ are colored orange, magenta, cyan, salmon, violet, light orange, teal, light pink, and lime green, respectively. **E**-**G**. the distance of N-terminal P12 and C-terminal E581 of Aα were shown, respectively. **A**-**I**. ENSA (this study, PDB ID 9XGY, under release), tpARPP19 (PDB ID 8TTB), FAM122A (PDB ID 8SO0), Eya3 (PDB ID 9C7T), p107 (PDB ID 9C6B), Eya3 (PDB ID 9N0Y), IER5 (PDB ID 8UO5), B55i (PDB ID 9N0Z), and apo PP2A(B55) (PDB ID 9MZW).

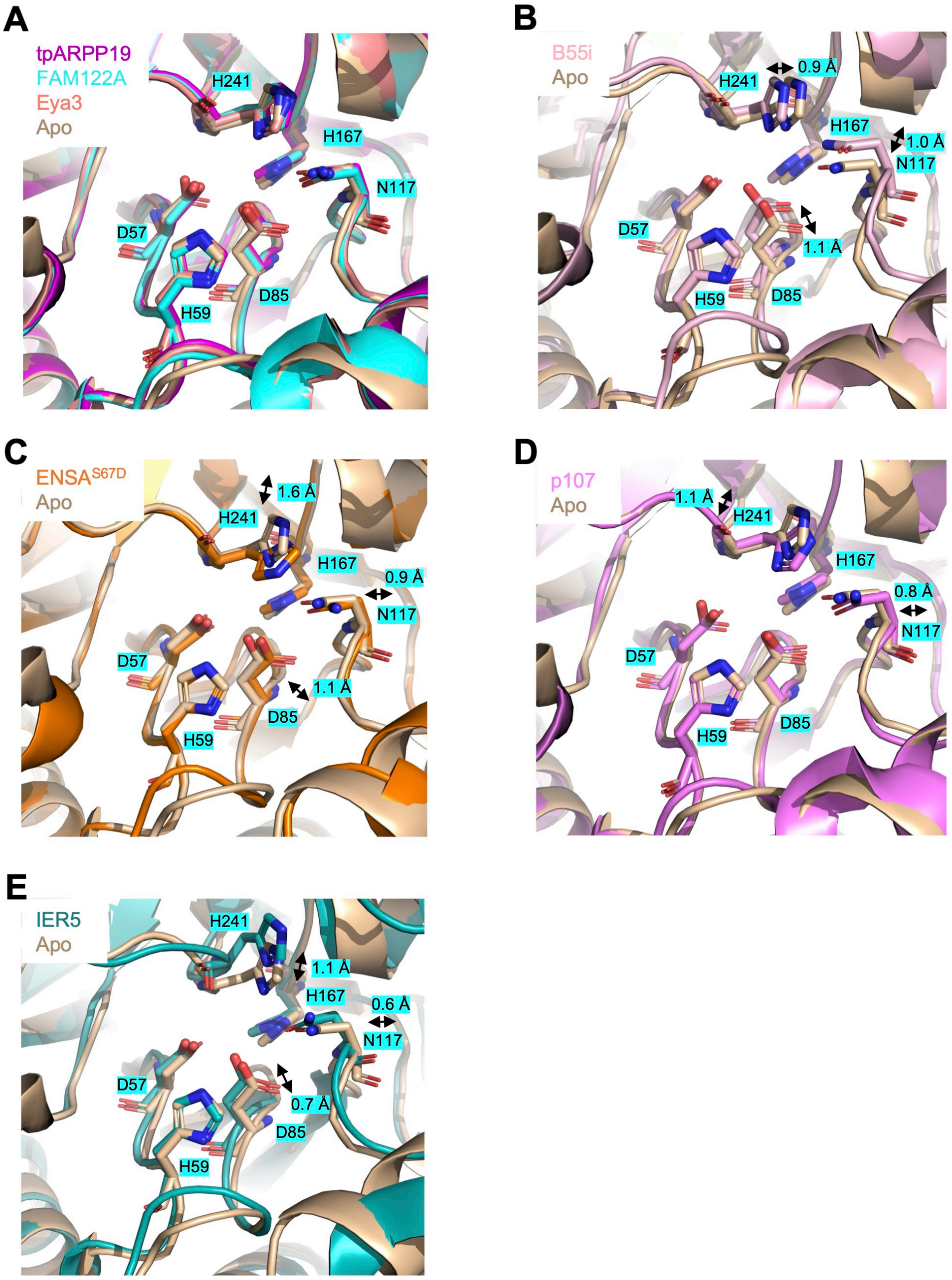


**Figure S12. Conformational changes of the PP2A-Cα active site residues upon inhibitors/recruiters/substrates binding.**

PP2A(B55) with differ inhibitors/recruiters/substrates are superimposed to Cα (residues 2-289) of apo PP2A(B55) (PDB ID 9MZW, wheat), respectively. The active site residues of C subunit are shown as sticks. **A**. tpARPP19 (PDB ID 8TTB, magenta), FAM122A (PDB ID 8SO0, cyan), and Eya3 (PDB ID 9C7T, salmon). **B**. B55i (PDB ID 9N0Z, light pink). **C**. ENSA$^{S67D}$ (this study, PDB ID 9XGY, under release, orange). **D**. p107 (PDB ID 9C6B, violet). **E**. IER5 (PDB ID 8UO5, teal).

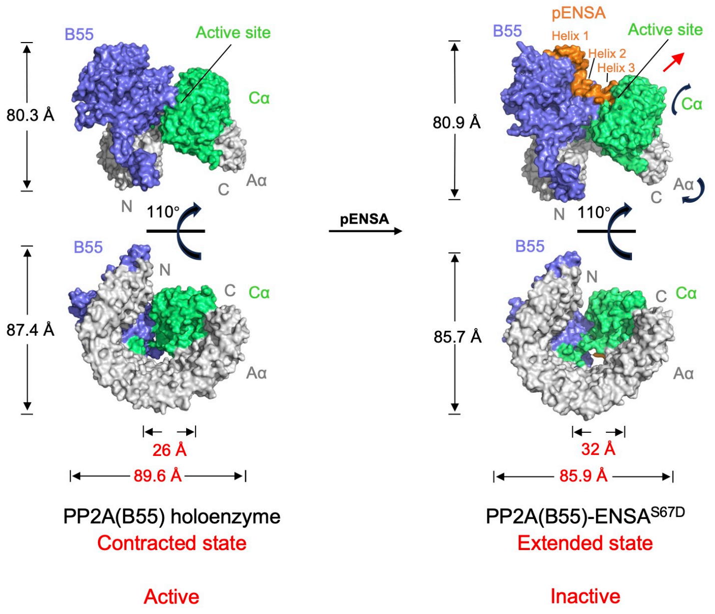


**Figure S13. The inhibit mechanism of PP2A(B55) holoenzyme by ENSA.** The schematic model proposes that the PP2A(B55α) holoenzyme in two distinct states: (1) An "Contracted state" active state when pENSA not bind. (2) A "Extended state" inactive state achieved when pENSA lock the pS67-containing motif into the catalytic active site.

**Supplementary Table 1. The binding kinetics parameters between $ENSA^{WT/S67D}$ and PP2A(B55) holoenzyme/subunits.**

| | | $K_D$ (μM) | $k_{on}$ ($M^{-1}*s^{-1}$) | $k_{off}$ ($s^{-1}$) | $R^2$ | |
|---|---|---|---|---|---|---|
| GST-$ENSA^{S67D}$ | PP2A (B55) | 0.66 ± 0.02 | 1.62E+05 | 1.07E-01 | 0.99 | |
| GST-$ENSA^{WT}$ | PP2A (B55) | 1.45 ± 0.13 | 2.14E+05 | 3.10E-01 | 0.99 | |
| GST-$ENSA^{S67D}$ | B55α | 17.2 ± 2.85 | 4.28E+04 | 7.35E-01 | 0.99 | |
| GST-$ENSA^{WT}$ | B55α | 4.95 ± 1.47 | 1.84E+05 | 9.13E-01 | 0.98 | |
| GST-$ENSA^{S67D}$ | Aα-Cα | N.R. | N.R. | N.R. | 0 | Double Ref |
| GST-$ENSA^{WT}$ | Aα-Cα | N.R. | N.R. | N.R. | 0 | Double Ref |
| GST-$ENSA^{S67D}$ | Cα | N.R. | N.R. | N.R. | 0.88 | Double Ref |
| GST-$ENSA^{WT}$ | Cα | N.R. | N.R. | N.R. | 0 | Double Ref |
| GST-$ENSA^{S67D}$ | Aα | N.R. | N.R. | N.R. | 0 | Double Ref |
| GST-$ENSA^{WT}$ | Aα | N.R. | N.R. | N.R. | 0 | Double Ref |

Double Ref means the BLI assay were measured by double referencing assay. N.R., no binding.

**Supplementary Table 2. The cryo-EM data collection and structure determination.**

| **Name** | **PP2A(B55)-ENSA$^{S67D}$** |
|---|---|
| EMDB and PDB ID (Under release) | (EMDB: EMD-66862) (PDB: 9XGY) |
| **Data collection and processing** | |
| Symmetry imposed | C1 |
| Final particle images (no.) | 227,045 |
| Map resolution (Å) | 3.03 |
| FSC threshold | FSC = 0.143 |
| Map resolution range (Å) | 4.50 to 2.5 |
| **Refinement** | |
| Initial model used (PDB) | 3DW8 and 2IAE |
| Model resolution (Å) | 3.28 |
| FSC threshold | FSC = 0.5 |
| Map sharpening B factor ($Å^2$) | -145.9 |
| Model composition | |
| Non-hydrogen atoms | 10978 |
| Protein residues | 1373 |
| Ligands | MN:2 |
| B factors ($Å^2$) | |
| Protein | 70.14 |
| Ligand | 102.95 |
| R.m.s. deviations | |
| Bond lengths (Å) | 0.003 |
| Bond angles (°) | 0.521 |
| Validation | |
| MolProbity score | 1.78 |
| Clashscore | 6.68 |
| Poor rotamers (%) | 1.97 |
| Ramachandran plot | |
| Favored (%) | 96.84 |
| Allowed (%) | 3.16 |
| Disallowed (%) | 0 |
